# CTHRC1 is a new therapeutic target and serum diagnostic biomarker for aortic dissection

**DOI:** 10.1101/2025.04.19.649636

**Authors:** Jiaxin Chen, Luzhang Ji, Jing Lu, Peng Wei, Yihao Li, Hui Huang, Chi Chen, Qiyong Li, Xin Dong, Tao Yu, Jin Tan, Guofang Xia, Jing Lin, Xianjing Wei, Ying Zhang, Liang Liu, Hao Huang, Cong Lu, Lianbin Wen, Yan Xiong, Zhongjing Cao, Jie Zhou, Tiantian Luo, Yu Chen, Peng Luo, Yang Chen, Jiayue Feng, Jinsong Li, Liang Wen, Lu Li, Xinyue Zhang, Guangre Xu, Chenzhang Ji, Huiqing Wan, Zhifeng Dong, Chengxin Shen, Congfeng Xu, Zhong Wu, Jin Zhou, Chenfei Wang, Wenjie Tian, Xiaoqing Wang

## Abstract

Aortic dissection (AD) is a cardiovascular disease with rapid onset and extremely high short-term mortality, currently lacking specific peripheral blood-based biomarkers and effective treatments. Here, our analysis of AD samples from animal models and human patients revealed elevated blood levels of CTHRC1, a protein secreted by vascular fibroblasts. Furthermore, CTHRC1 regulates the phenotypic switch of vascular smooth muscle cells (VSMCs) during arterial remodeling by binding to ADAM9 on the VSMC membrane, activating the ERK1/2 signaling pathway, and promoting a contractile-to-synthetic transition. In Ang-II/BAPN induced mouse models, genetic ablation or antibody-mediated blockade of CTHRC1 effectively prevented AD development. These findings unveil CTHRC1 as a critical regulator of VSMC phenotype and aortic structural integrity via the ERK1/2 pathway, suggesting its potential as a novel serum diagnostic biomarker for AD diagnostic and a promising therapeutic target.

**Graphical abstract:** 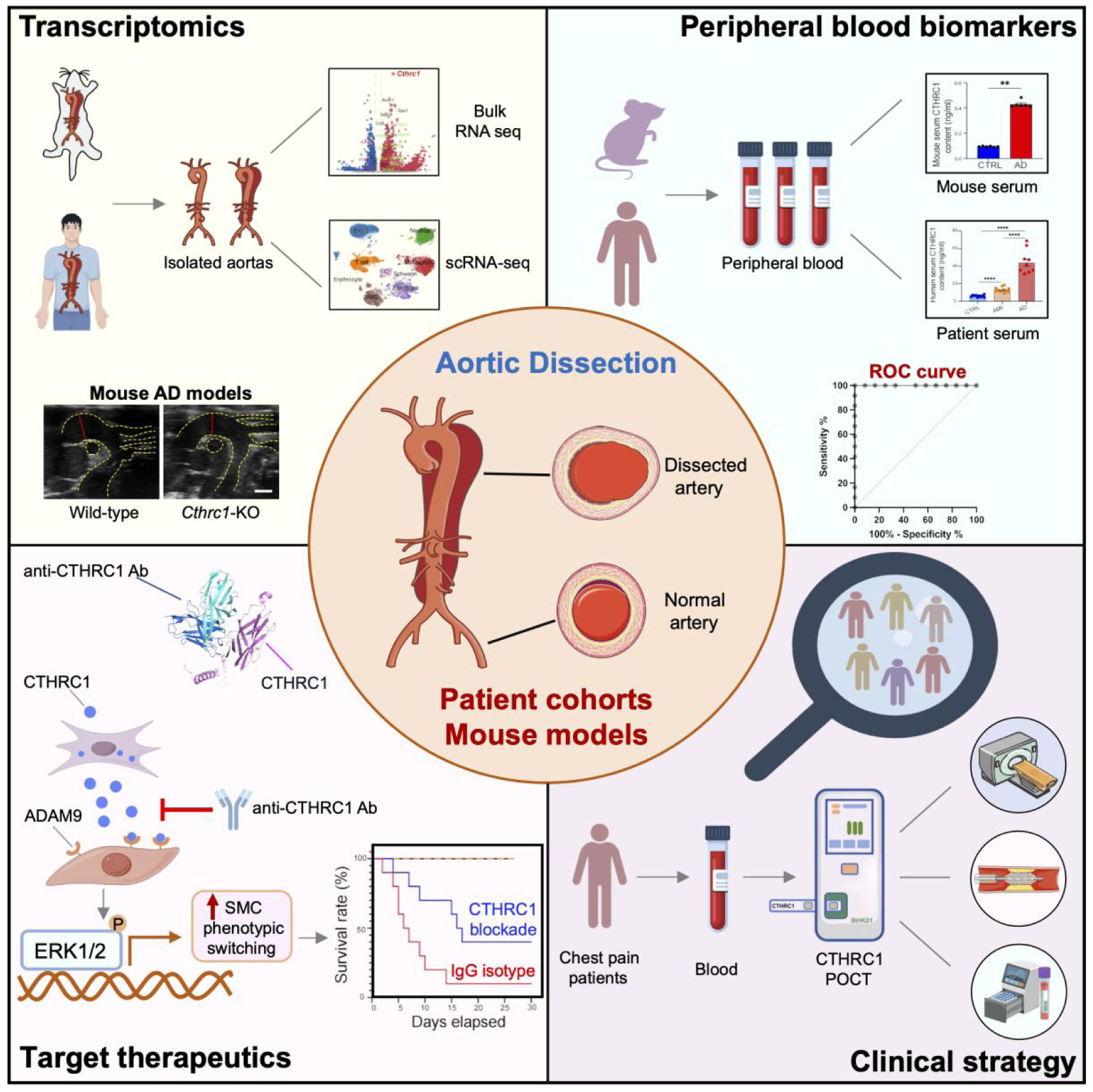

**In brief:** CTHRC1 is aberrantly expressed in aortic adventitial fibroblasts from dissected aortas and function as an exocrine mediator that induces phenotypic switching of vascular smooth muscle cells. A monoclonal antibody targeting CTHRC1 demonstrates therapeutic potential in mouse models of aortic dissection.

**Highlights:**

- CTHRC1 is highly expressed in aortic adventitial fibroblasts from dissected aortas
- CTHRC1 interacts with ADAM9 to induce phenotypic switching of VSMCs
- An anti-CTHRC1 antibody inhibits aortic dissection formation*invivo*
- Elevated levels of serum CTHRC1 can be used to identify AD in patients presenting with chest pain

## Introduction

Aortic dissection (AD) is a devastating cardiovascular characterized by acute onset, rapid progression, and a high short-term mortality rate.^1-2^ In response to mechanical and pathological stress, arterial remodeling usually occurs, destabilizing the aortic wall integrity and rendering it vulnerable to enlargement.^3-6^ AD is defined by a tear in the intima and the formation of a false lumen within the aortic wall layers, which can ultimately penetrate the adventitia and lead to uncontrolled hemorrhage.^7,8^ Population-based studies indicate that the incidence and mortality rates of AD have gradually increased over recent decades, largely due to limited diagnostic and therapeutic strategies.^9-11^

The aortic wall is composed of three layers, which are the tunica intima, the tunica media, and the tunica adventitia. These layers are predominantly composed of endothelial cells,^12,13^ vascular smooth muscle cells (VSMCs),^14-16^ fibroblasts, immune cells^17^, and extracellular matrix components.^18^ These cellular and extracellular components perform sophisticated and coordinated functions to withstand dynamic changes during the process of mechanical stress, injury, repair, and remodeling. The endothelial tight junction-based barrier located in the intima plays a critical role in the development of thoracic aortic aneurysm and dissection, suggesting a potential mechanism underlying this disease.^19^ Recent studies have highlighted the heterogeneity of VSMCs, revealing important findings regarding distinct VSMC subpopulations and their phenotypic transitions during the formation of aortic aneurysms and dissections, particularly through single-cell RNA sequencing.^20^ Maintaining VSMC contractile phenotype and preventing its transformation into a synthetic state presents new opportunities for intervening in the progression of AD.^21^ Additionally a CARTPT-expressing VSMC subtype identified has been associated with medial calcification of the aorta.^22^ Overall, dysfunction in endothelial cells and VSMCs likely contributes to aortic wall integrity deterioration, leading to AD.

Recently, there has been a growing appreciation of the importance of the outermost layer of the aorta to vascular biology.^23-25^ Several lines of compelling evidence have illuminated the role of resident fibroblasts in depositing collagen fibrils and extracellular matrix components in the vasculature.^26,27^ Under pathological conditions, activated fibroblasts, commonly referred to as myofibroblast,^28-30^ enhanced the inflammatory response during the development of AD.^31,32^ Despite ongoing research aimed at elucidating the role of fibroblasts in stressed aortic tissue,^33,34^ the specific cell-cell interactions of fibroblasts with other components of the aorta underlying the pathogenesis of AD remain poorly understood.

Here, we constructed a single-cell transcriptomic landscape of dissected aortas from both mouse models and patients with AD. This analysis uncovered a new fibroblast subpopulation characterized by high expression of *CTHRC1*. Using genetic ablation of *Cthrc1* in mouse models and anti-CTHRC1 monoclonal antibodies, we provide evidence for the critical role of this specific fibroblast population in the pathogenesis of AD. Blockade of *Cthrc1* inhibited the transition of VSMCs from a contractile to a synthetic phenotype, ameliorating the progression of AD and improving survival rates *in vivo*. Further functional analyses demonstrated that adventitial fibroblast-derived exocrine CTHRC1 interacts with ADAM9 on VSMCs, activating the downstream ERK1/2 signaling pathway. Thus, the CTHRC1-ADAM9-ERK1/2 axis may represent a critical signaling pathway that promotes detrimental VSMC phenotypic switching. We also identified significantly elevated levels of serum CTHRC1 in both AD mouse models and human patients. Our findings suggest that serum CTHRC1 can effectively differentiate AD patients from those with acute myocardial infarction (AMI). Overall, this study highlights the unique role of CTHRC1 in vascular remodeling, supporting its potential as a diagnostic biomarker and therapeutic target for AD in future clinical applications.

## Results

### CTHRC1 is highly expressed in the aortas of mouse models and patients with AD

Mice of different genders infused with angiotensin II (Ang-II) and β-aminopropionitrile (BAPN) using mini-osmotic pumps are widely recognized as a reliable model for studying AD.^35-37^ In this study, we first collected aortas from C57BL/6 mice infused with PBS or Ang-II/BAPN for 14 days and performed bulk RNA-seq to acquire comprehensive transcriptomic profiles. Differential expression analysis identified 4,049 up-regulated genes and 1,258 down-regulated genes, with *Cthrc1* showing significant upregulation in AD mouse models (Figure S1A). Additionally, previously reported AD-associated genes, such as *Lgmn*,^38^ *Hif1a*,^39^ *Slc44a5,*^40^ *Hmox1*,^41^ and *Tyrobp*,^42^ were significantly elevated. Meanwhile, matrix metalloproteinases (MMPs) like *Mmp2* and *Mmp9*, which are involved in vascular remodeling and extracellular matrix (ECM) degradation,^43,44^ were also significantly upregulated in the aortas of AD mice. KEGG enrichment analysis of differentially expressed genes (DEGs) indicated that the up-regulated genes were enriched in the MAPK signaling pathway, PI3K-Akt signaling pathway, and IL−17 signaling pathway, while the down-regulated genes were enriched in vascular smooth muscle contraction, cAMP signaling pathway, and regulation of the actin cytoskeleton (Figure S1B). These signaling pathways have been previously linked to the modulation of AD.

To further locate the AD-associated genes within the tissue microenvironment, aortas were processed separately and subjected to single-cell RNA sequencing (scRNA-seq) to investigate the heterogeneity of aortic cell clusters (Figure 1A). After filtering out low-quality cells (Figure S1C), we performed integrative unsupervised cell clustering analysis using Scanpy^45^ and identified 15 major cell clusters (Figure S1D). Guided by representative cell type-specific markers,^46^ we annotated endothelial cells (ECs), smooth muscle cells (SMCs), fibroblasts (FBs), immune cells, and neuron-related cells (Figures 1B and S1E). We compared the dynamics of cell clusters between the control and AD groups and observed a smaller SMC cluster in AD samples, while the fibroblast cluster was significantly increased (Figure 1C). To further characterize the tunica adventitia, we subdivided fibroblast clusters into 2 groups (FB1 and FB2) and identified expression markers for each subcluster (Figures 1D and S1F). Notably, *Cthrc1* was highly expressed in the FB1 subcluster but not in FB2. Furthermore, we found that FB1 cells were predominantly derived from AD mouse models, whereas FB2 consisted mainly of cells originating from normal aortas (Figure 1E). These data suggest that *Cthrc1*^+^ fibroblasts may represent an AD-specific emerging fibroblast cluster.

**Figure 1.**
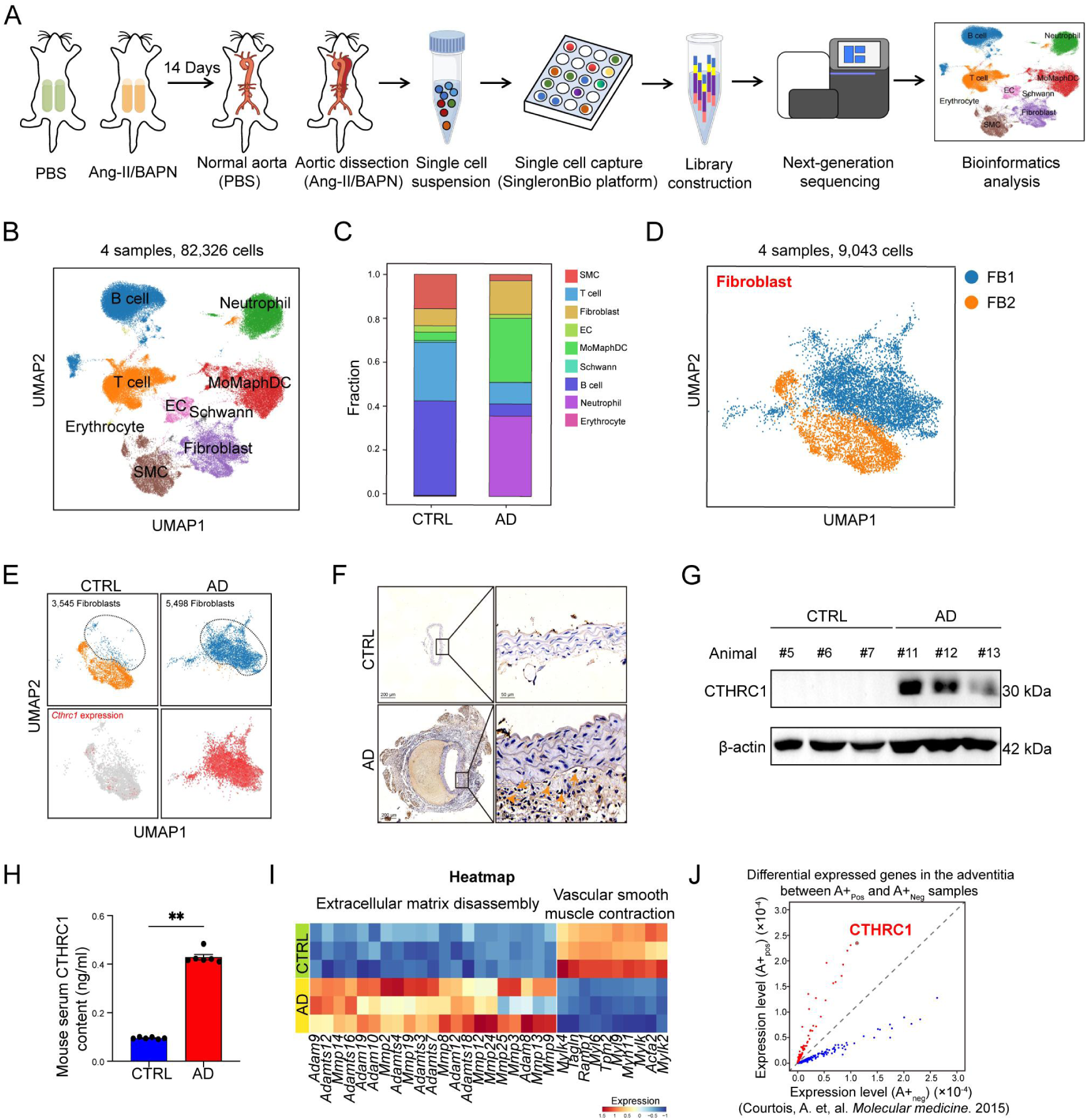
CTHRC1 is highly expressed in the aortas of mouse models and patients with AD. (A) A schematic diagram illustrating the experiment design. Single-cell RNA sequencing was applied to 4 samples from C57BL/6 mice that were infused with saline or simultaneously administrated by Ang-II/BAPN for 14 days. (B) UMAP plot of the 82,326 single cells from the 4 samples outlined in (A), showing the identification of 9 main clusters. (C) The proportion of 9 cell clusters in control and AD mice. (D) UMAP plot of the 9,043 fibroblasts with 2 subclusters. (E) UMAP plots showing comparison of fibroblast clusters or Cthrc1 expression between the AD and control group. (F) Immunohistochemical staining for CTHRC1 in mouse aortas. The orange arrows indicating CTHRC1 expression in adventitial fibroblasts of the dissected aortas. (G) Representative images showing immunoblotting analysis of CTHRC1 levels in control and AD mice (n=6). (H) Mouse serum CTHRC1 content in control and AD groups (n=6). Data are presented as mean ± SEM. (Student’s t test, \*\**P* ≤ 0.01.) (I) Heatmap of expression levels of genes related to vascular smooth muscle cell contraction and extracellular matrix disassembly between control and AD mice. (J) Scatterplot showing gene expression level in the adventitia between A+_Pos_ and A+_Neg_ samples. Red: upregulated genes in the adventitia of aortas from patients prone to aorta rupture, blue: downregulated genes. See also Figure S1.

Next, we collected aortic tissues and peripheral blood from control and AD mice for further analysis. Immunohistochemistry (IHC) staining revealed that CTHRC1 protein was predominantly localized to adventitial fibroblasts in the dissected aortas (Figure 1F). Immunoblotting and serum ELISA tests also demonstrated significantly increased CTHRC1 protein levels in AD mice compared to controls (Figures 1G and 1H). Moreover, an upregulation of ECM-related metalloproteinases and a downregulation of VSMC contractile proteins were observed during AD formation, as illustrated by heatmaps (Figure 1I). To explore whether CTHRC1 contributes to the pathogenesis of AD in humans, we reanalyzed publicly available datasets.^21,47^ We found that, compared to control individuals, the expression level of *CTHRC1* was markedly up-regulated in the adventitia of patients prone to aorta rupture (Figure 1J) and in the fibroblasts of aortas from cohorts with thoracic aortic aneurysms (Figure S1G). At the level of cell lines, we found that the fibroblast cell line NIH-3T3 showed a dramatic increase in CTHRC1 protein level after Ang-II/BAPN treatment, while no such increase was observed in endothelial HUVECs or smooth muscle cell line MOVAS (Figures S1H-S1J). Although several studies have noted potential gender differences in the presentation and outcomes of AD,^9,48,49^ the consistency between male and female mice infused with PBS or Ang-II/BAPN was evaluated at both the gene and protein levels (Figures S1K-S1O). In female AD mice, similar to males, elevated CTHRC1 expression was observed in response to pathological stimuli. Taken together, these data suggest an essential role for CTHRC1 in the formation of AD.

### *Cthrc1* KO attenuates AD formation in mouse models

To gain insight into the function of CTHRC1, we generated genetic knockout (*Cthrc1^-/-^*) C57BL/6J mice by deleting *Cthrc1* exons 2 and 3. Six-week-old male *Cthrc1^-/-^* mice and wide-type (WT) mice were treated with PBS or Ang-II/BAPN for 14 days (Figure 2A). In *Cthrc1^-/-^* mice, Cthrc1 was successfully deleted in the aortas, regardless of whether they were infused with PBS or Ang-II/BAPN (Figures 2B and S2A). Ultrasound imaging demonstrated a remarkable decrease in the maximal aortic diameter of *Cthrc1^-/-^* mice administrated with Ang-II/BAPN on day 14 compared to WT controls (Figures 2C and 2D). Consistently, the incidence of AD in *Cthrc1^-/-^* mice was significantly lower than in WT mice. Both ascending and abdominal aortas exhibited a notably decreased occurrence of AD in *Cthrc1^-/-^* mice (Figures 2E and 2F). Kaplan-Meier survival analysis indicated a significantly higher survival rate for *Cthrc1^-/-^* mice challenged with Ang-II/BAPN compared to WT mice (37.5 vs. 87.5%) (Figure 2G). Furthermore, Hematoxylin and eosin (H&E) and Elastica van Gieson (EVG) stainings revealed decreased aortic diameters and fewer elastin breaks in *Cthrc1^-/-^* mice compared to WT (Figures 2H and 2I). Additionally, aortas from *Cthrc1^-/-^* mice showed lower levels of mucoid extracellular matrix accumulation than WT controls (Figure S2B). Collectively, these data indicate a beneficial effect of *Cthrc1* deficiency in the formation and progression of AD.

**Figure 2.**
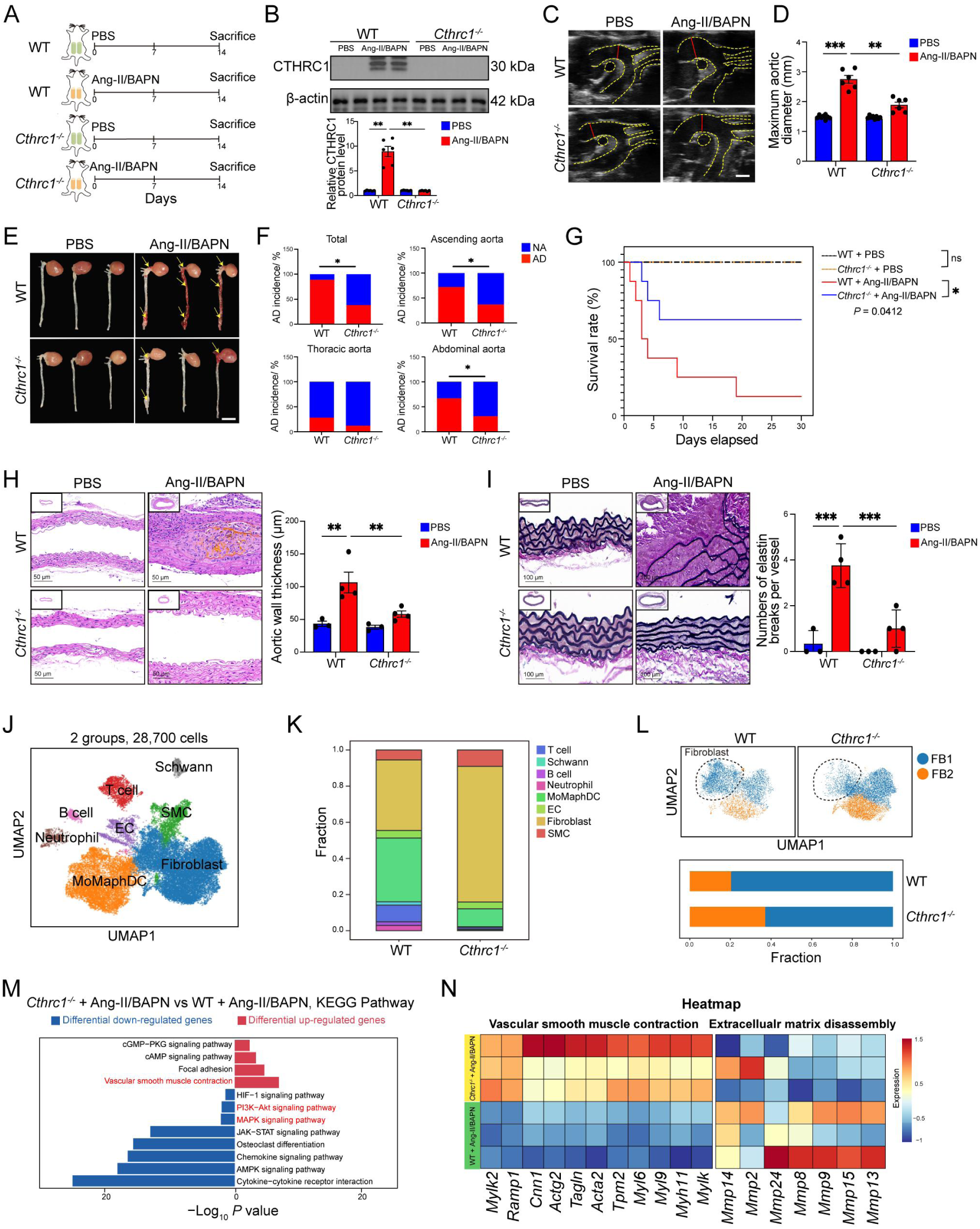
*Cthrc1* KO attenuates AD formation in mouse models. (A) Experiment design. 6-week-old male *Cthrc1^-/-^* mice (n=16) and wide type (WT) mice (n=18) were treated with saline or Ang-II/BAPN for 14 days. (B) Representative images of immunoblotting analysis of CTHRC1 levels in aortas from the mice in A. Bottom: quantification of CTHRC1 expression in aortas. (a ordinary one-way ANOVA followed by Bonferroni’s multiple comparison test, \*\**P* ≤ 0.01.) (C) Representative ultrasound images of aorta in mice (scale bar, 1mm). (D) Measurements of maximum aortic diameter of mice in A (n=6-9 per group). (a ordinary one-way ANOVA followed by Bonferroni’s multiple comparison test, \*\**P* ≤ 0.01, \*\*\**P* ≤ 0.0001.) (E) Representative pictures of whole aortas from the mice in A. The yellow arrows indicate the dissected sites of vascular lesions. (F) Quantification of AD incidence in the whole aorta, ascending, thoracic and abdominal aortas from WT and *Cthrc1^-/-^* mice. (Fisher’s exact test, \**P* ≤ 0.05.) (G) Kaplan-Meier survival rates analysis of the mice in A. (Kaplan-Meier survival rates analysis, \**P* ≤ 0.05, ns, no significance.) (H) Representative images of histological staining with hematoxylin and eosin in aortic sections from the mice in A. Right: measurements of aortic wall thickness of aortas (scale bar, 50 μm). (a ordinary one-way ANOVA followed by Bonferroni’s multiple comparison test, \*\**P* ≤ 0.01.) (I) Representative images of Elastica van Gieson staining in aortic sections from the mice in A. Right: measurement of numbers of elastin breaks per vessel (scale bar, 100 μm). (a ordinary one-way ANOVA followed by Bonferroni’s multiple comparison test, \*\*\**P* ≤ 0.0001). (J) UMAP plot of the 28,700 single cells from WT and *Cthrc1^-/-^* mice challenged with Ang-II/BAPN for 14 days, displaying the identification of 8 major clusters. (K) The proportion of 8 cell clusters in WT and *Cthrc1^-/-^* mice infused with Ang-II/BAPN. (L) Upper: UMAP plot of fibroblast subclusters in the WT group and *Cthrc1^-/-^* group respectively. Bottom: the proportion of FB1 and FB2. (M) KEGG enrichment analysis of differentially expressed genes in aortic tissues from WT and *Cthrc1^-/-^* mice analyzed by bulk-RNA seq data. (N) Heatmap showing the expression levels of genes related to vascular smooth muscle cell contraction and extracellular matrix disassembly. See also Figure S2.

To clarify the response of different cell populations affected by CTHRC1, we performed scRNA-seq using the aortas of mice challenged with Ang-II/BAPN. By analyzing cell-type-specific markers from 28,700 high-quality cells, we identified 8 major cell clusters, including ECs, SMCs, fibroblasts, MoMaphDCs, Schwann cells, T cells, B cells, and neutrophils (Figure 2J). Cell distribution analysis showed that knockout of *Cthrc1* altered the general proportion of major cell clusters, such as SMCs, fibroblasts, MoMaphDCs, and neutrophils (Figure 2K). Notably, the proportion of harmful FB1 subcluster was decreased upon *Cthrc1* deficiency, compared to the WT controls (Figure 2L). We also performed bulk RNA-seq of aortas from AD-induced WT and *Cthrc1^-/-^* mice. Genes that showed decreased expression levels upon *Cthrc1* deficiency were associated with the MAPK and PI3K-Akt signaling pathways, as well as extracellular matrix disassembly (e.g., *Mmp2*, *Mmp9*, and *Mmp15*). In contrast, upregulated genes were involved in vascular smooth muscle contraction (e.g., *Myh11*, *Tagln*, and *Cnn1*) (Figures 2M, 2N, S2C, and S2D). Moreover, metalloproteinases, ADAMs (a disintegrin and metalloproteinases), and ADAMTSs (ADAMs with a thrombospondin motif), known to participate in the turnover of ECM proteins for vascular remodeling, were also affected.^50-55^ Specifically, family members such as *Adamts16*, *Adamts19*, *Adam12*, and *Adam10* were downregulated in the aortas of the *Cthrc1^-/-^* AD group (Figures S2E and S2F). It is widely acknowledged that men are more commonly affected by AD than women and that women tend to develop AD later in life.^56,57^ Our data indeed showed that female WT mice exhibited a lower incidence of AD formation than males. Nevertheless, the loss of *Cthrc1* further mitigates the formation of AD in female mice (Figures S2G-S2Q). These alterations may also be partially due to the activation of VSMC contractile genes, including *Acta2*, *Tagln*, and *Mylk* (Figures S2R and S2S). In summary, these findings indicate that the knockout of *Cthrc1* abrogates the phenotypic switching of VSMCs from contractile to synthetic states, reinforcing our understanding of its pivotal role in AD progression.

### Systematic transcriptome analysis reveals communication between fibroblasts and VSMCs

The vascular microenvironment is a complex and dynamic mixture of numerous components, including fibroblasts, VSMCs, immune cells, ECs, and soluble factors.^58^ To investigate whether CTHRC1 affects cell-cell communications (CCC) during AD progression, we constructed CCC networks among all major cell types using CellChat.^59^ Overall, the number of inferred interactions and the strength of these interactions were substantially increased in the dissected aortas from AD models compared to controls (Figure 3A). Furthermore, fibroblast-SMC interactions were enhanced during AD development (Figure 3B). Next, we explored the signaling patterns involved in outgoing interactions between fibroblasts and SMCs, identifying Spp1 signaling,^27,60^ Fn1 signaling, Pros1-Mertk signaling, and Gas6-Mertk signaling as highly enriched and activated in AD (Figure 3C). These results indicate that fibroblasts play a significant role in signaling interactions with SMCs during AD progression, highlighting a novel fibroblast-mediated signaling mechanism in vascular remodeling.

**Figure 3.**
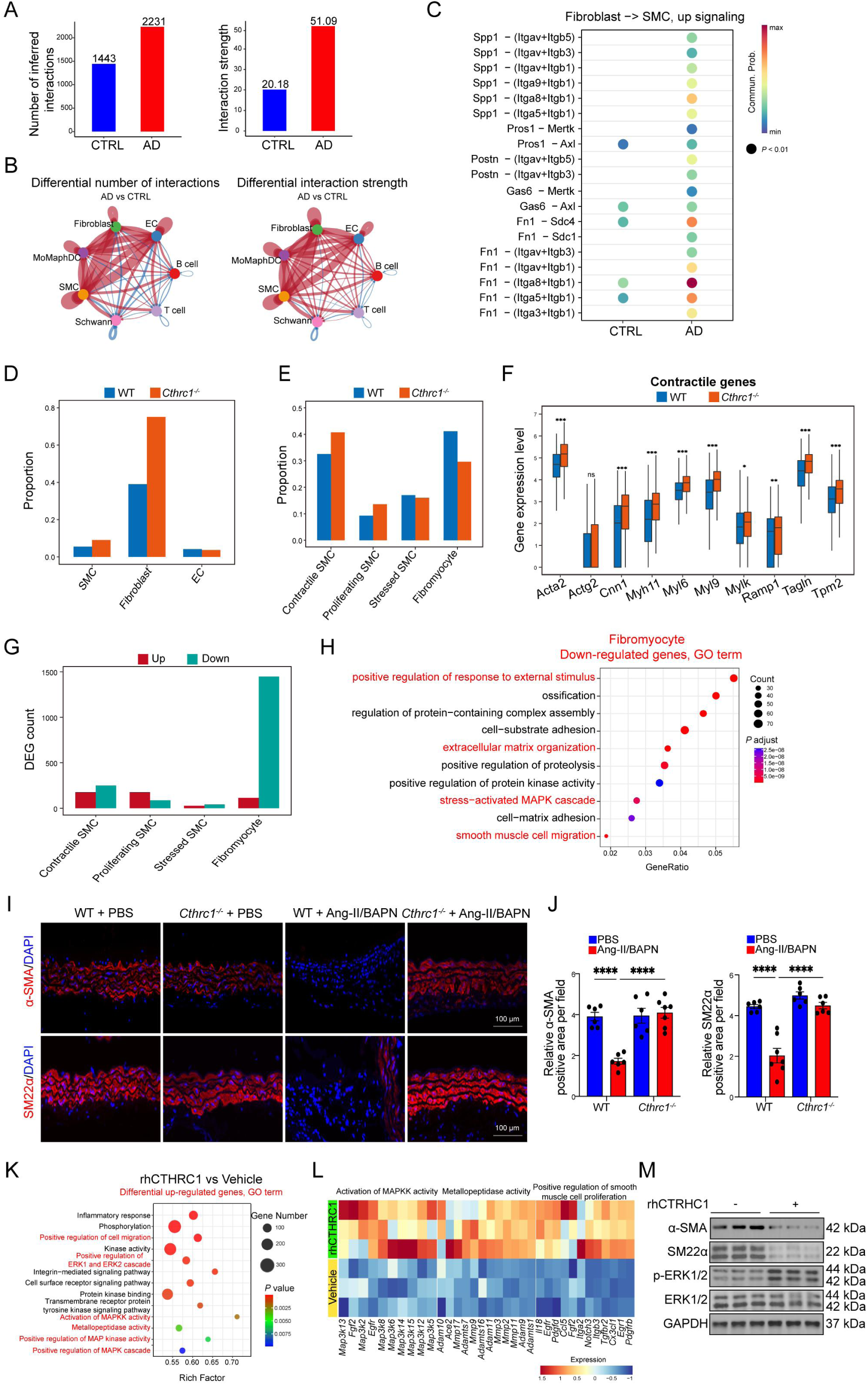
Systematic transcriptome analysis reveals communication between fibroblasts and VSMCs. (A) Comparison of the total number of cell-cell interactions and the interaction strength from the scRNA-seq data of control and AD mice. (B) Circle plot showing differential cell-cell interaction numbers and strength between fibroblast, SMC, EC, MoMaphDC, Schwann, B cell and T cell clusters predicted by CellChat. Each circle represents one cell cluster, edges between circles represent intercellular signaling between cell clusters, and edge thickness reflects interaction number or strength. Red edges represent increased interaction number or strength and blue represent decreased interaction number or strength in AD mice versus control mice. (C) Identification of up-regulated signaling by comparing the communication probabilities mediated by ligand–receptor pairs from fibroblasts to SMCs in AD mouse models. (D) D to H: scRNA-seq of the two aortas from WT and *Cthrc1^-/-^* mice infused with Ang-II/BAPN for 14 days were pooled per group. The proportion of major cell types including SMCs, ECs and FBs in Ang-II/BAPN-infused WT and *Cthrc1^-/-^* mice. (E) Bar diagram showing the percentage of 4 subclusters in aortic SMCs from the two groups of mice. (F) Boxplot illustrating the differential gene expression of contractile genes in aortic SMCs of the two mouse groups. (\**P* ≤ 0.05, \*\**P* ≤ 0.01, \*\*\**P* ≤ 0.0001. ns, no significance.) (G) Barplot depicting the numbers of differential expressed genes in SMC subclusters of the two mouse groups. (H) Gene Ontology enrichment analysis of down-regulated genes in fibromyocytes of the two mouse groups. (I) Immunofluorescence staining of α-SMA and SM22α expression in aortas from Ang-II/BAPN-infused WT and *Cthrc1^-/-^* mice. DAPI: 4’,6-diamidino-2-phenylindole (scale bar: 100 μm). (J) Quantification of α-SMA and SM22α positive area in aortas. (a ordinary one-way ANOVA followed by Bonferroni’s multiple comparison test, \*\*\*\**P* ≤ 0.00001.) (K) K to L: Incubating the murine aortic smooth muscle cells (MOVAS) with rhCTHRC1 in vitro. Gene Ontology enrichment analysis of up-regulated genes in MOVAS cells treated with rhCTHRC1 by bulk RNA-seq. (L) Heatmap showing the expression levels of genes related to metallopeptidase activity, activation of MAPKK activity and positive regulation of smooth muscle cell proliferation in MOVAS cells treated with rhCTHRC1 or Vehicle. (M) Immunoblotting analysis of the protein levels of α-SMA, SM22α, and p-ERK1/2 in MOVAS with or without rhCTHRC1 treatment. See also Figure S3.

The dominant role of *Cthrc1^+^* fibroblasts in AD development, coupled with their strong intercellular communications with SMCs, prompted us to further investigate the specific functions of CTHRC1 in SMC phenotypic modulation. Our scRNA-seq data of aortas (Figures 2 and S3A-S3D) showed that the proportion of SMCs, a major cell type in the tunica media, increased upon *Cthrc1* knockout (Figure 3D). We then tried to explore whether *Cthrc1* deficiency influences SMC subtypes, identifying four SMC subclusters: contractile SMCs, proliferating SMCs, stressed SMCs, and fibromyocytes based on representative marker genes (Figures S3E and S3F).^46^ Compared to controls, *Cthrc1^-/-^* mice exhibited partial preservation of the reduction in contractile SMCs during AD development (Figure 3E). In contrast, the proportion of fibromyoctes among all SMCs from *Cthrc1^-/-^* mouse aortas was lower than in controls (Figure 3E). Overall, contractile genes^61,62^ showed increased expression levels in SMCs from *Cthrc1^-/-^* mouse samples (Figure 3F). We further identified DEGs for each SMC subcluster and found that fibromyocytes exhibited the highest number of DEGs after the loss of *Cthrc1* (Figure 3G). Functional enrichment analysis revealed that down-regulated genes in fibromyocytes were related to positive regulation of response to external stimulus, stressed-activated MAPK cascade, and smooth muscle cell migration (Figure 3H), while up-regulated genes in contractile SMCs were involved in the muscle system process, striated muscle contraction, and skeletal muscle contraction (Figure S3G). Collectively, *Cthrc1^+^* fibroblasts drive the transition of SMCs from a contractile to a synthetic phenotype by altering cell signaling, contributing to unfavorable vascular remodeling during AD.

In light of our findings that *Cthrc1* inhibition abrogated the effect on contractile-to-synthetic phenotype switching *in vivo* at the transcriptional level, we further confirmed these results at the protein level both *in vivo* and *in vitro*. Immunofluorescence (IF) and immunoblotting analyses demonstrated the preservation of SMC contractility marker proteins SM22α and α-SMA in the aortas of *Cthrc1^-/-^* mice compared to WT mice exposed to Ang-II/BAPN (Figures 3I, 3J, S3H and S3I). Given that biologically active CTHRC1 is secreted from fibroblasts under pathological conditions (Figure 1H), we treated aortic smooth muscle cell, MOVAS^63^ with recombinant CTHRC1 (rhCTHRC1) and validated its effects *in vitro*. RNA-seq was performed to discern differentially expressed genes in MOVAS cells treated with rhCTHRC1 or vehicle (Figure S3J). Genes associated with the positive regulation of the Erk1 and Erk2 cascade, metallopeptidase activity, and positive regulation of the MAPK cascade were enriched in the 1,148 up-regulated genes in MOVAS cells treated with rhCTHRC1 (Figures 3K and 3L). Consistently, KEGG pathway enrichment analysis indicated that these up-regulated genes were related to the MAPK and PI3K-Akt signaling pathways (Figure S3K). At the protein level, we found that SMC contractility markers SM22α and α-SMA exhibited significantly lower expression levels, along with higher ERK1/2 phosphorylation, in the presence of rhCTHRC1 (Figure 3M). These findings consolidated the critical role of CTHRC1 in promoting VSMC differentiation through ERK1/2 phosphorylation in AD pathogenesis.

### CTHRC1 regulates phenotypic switching of VSMCs by binding with ADAM9

Next, we tried to identify the potential membrane-anchored protein responsible for CTHRC1’s stimulation of SMC phenotypic switching. We first incubated MOVAS cells with rhCTHRC1 and collected cell membrane lysates to perform co-immunoprecipitation (Co-IP) with anti-CTHRC1 antibodies, along with mass spectrometry (MS) to identify CTHRC1 binding proteins (Figures 4A and S4A). More than 18 membrane-anchored proteins were pulled down in two independent assays (Figure 4B). Among these candidates, ADAM metallopeptidase domain 9 (ADAM9) ranked as the most prominent (Figure S4B). Previous literature indicates that members of the ADAM family are associated not only with AD or aortic aneurysm,^43,50,64,65^ but also with the regulation of ERK1/2 signaling.^66-69^ Thus, we hypothesized that ADAM9 might serve as an important binding protein on VSMCs to facilitate signal transduction between CTHRC1 and ERK1/2, leading to CTHRC1-mediated SMC phenotypic switching. Co-IP combined with immunoblotting confirmed the physical interaction between CTHRC1 and ADAM9 (Figure 4C). Furthermore, rhCTHRC1 incubation dramatically upregulated the expression levels of ADAM9 in MOVAS cells (Figures S4C and S4D), consistent with our *in vivo* RNA-seq data (Figure S4E) and previous findings in aortas from Ang-II-challenged mice.^70^

**Figure 4.**
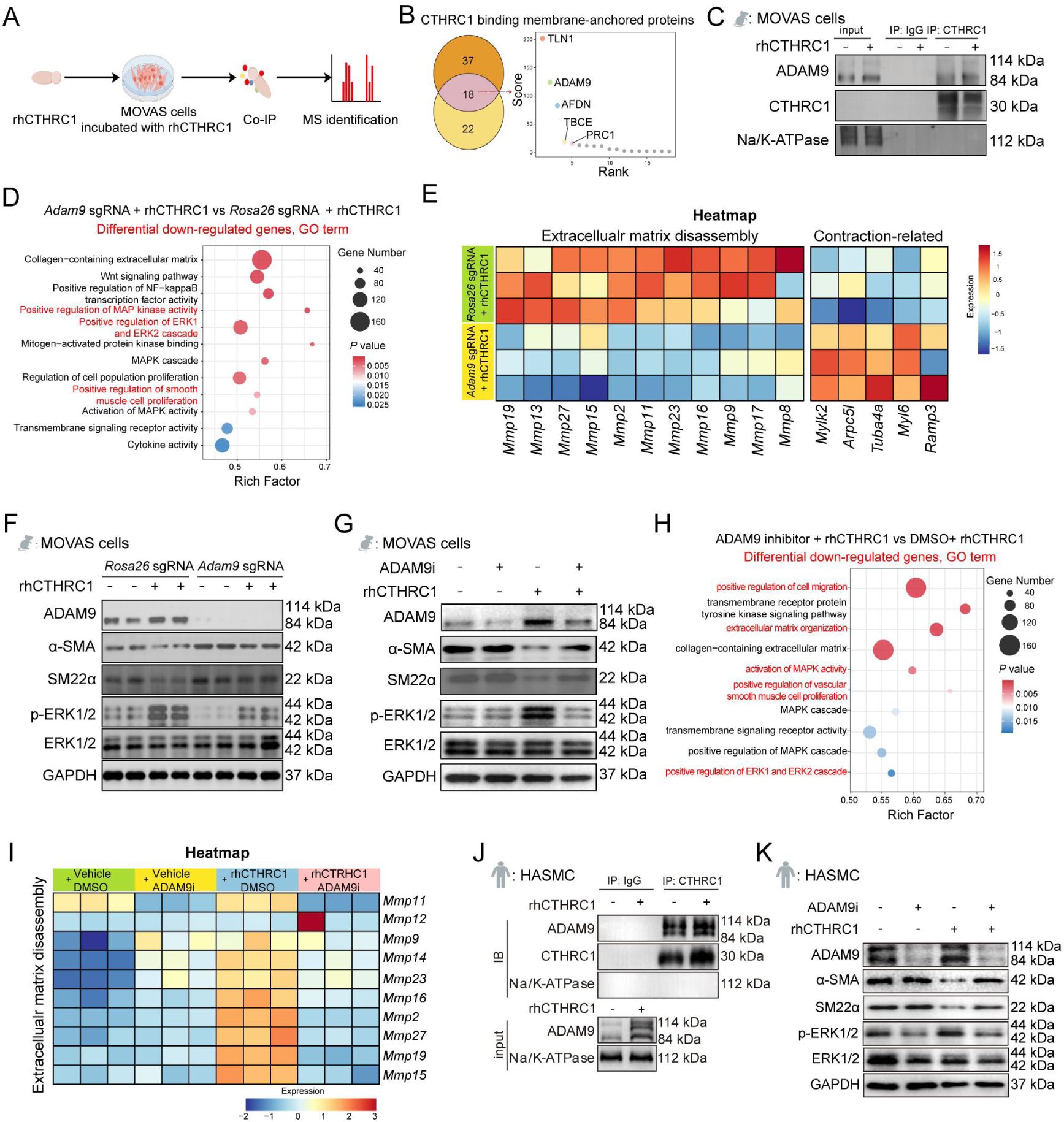
CTHRC1 regulates phenotypic switching of VSMCs by binding with ADAM9. (A) Workflow of the immunoprecipitation (IP) of CTHRC1 identified by mass spectrometry (MS). (B) Venn diagram of intersection analyses of two independent assays for potential receptors of CTHRC1 on MOVAS cells. Right: Scores of candidate binding proteins identified by MS. (C) Co-IP-blotting of CTHRC1 with ADAM9 receptor on MOVAS cells. (D) D to E: whole transcriptome analysis of MOVAS cells incubated with rhCTHRC1, knockouting *Roas26* (control) or *Adam9* (*Adam9*-KO). Gene Ontology enrichment analysis of down-regulated genes in *Adam9*-KO MOVAS cells treated with rhCTHRC1. (E) Heatmap showing the expression levels of genes associated with extracellular matrix disassembly and contraction. (F) Immunoblotting analysis of ADAM9, α-SMA, SM22α, and p-ERK1/2 expression in MOVAS cells, knockouting *Roas26* or *Adam9*. (G) Immunoblotting analysis of ADAM9, α-SMA, SM22α, and p-ERK1/2 expression in MOVAS cells with or without treatment of ADAM9 inhibitor (ADAM9i). (H) H to I: Comprehensive profiles of genes in MOVAS cells with treatment of ADAM9i analyzed by bulk RNA-seq. Gene Ontology enrichment analysis of down-regulated genes in MOVAS cells treated with ADAM9i. (I) Heatmap showing the expression levels of metalloprotein genes in MOVAS MOVAS cells in the presence of rhCTHRC1cells treated with ADAM9i or DMSO. (J) Interaction of CTHRC1 with ADAM9 in human aortic smooth muscle cells (HASMCs) confirmed by Co-IP-blotting. (K) Western blotting analysis of ADAM9, α-SMA, SM22α, and p-ERK1/2 expression in HASMCs with treatment of ADAM9i. See also Figure S4.

We then performed functional studies to examine the effects of the CTHRC1-ADMA9 interaction on VSMC phenotypic transformation and intracellular signal transduction. We conducted bulk RNA-seq and immunoblotting analysis using MOVAS cells infected with either *Rosa26* sgRNA (control) or *Adam9* sgRNA (*Adam9*-KO) lentivirus (Figures 4D to 4F; Figures S4F to S4H). Whole-transcriptome analysis revealed 1,245 differentially expressed genes between *Adam9*-KO and control cells incubated with rhCTHRC1 (Figure S4F). The down-regulated genes in *Adam9*-KO cells were functionally annotated for positive regulation of the ERK1 and ERK2 cascade, activation of MAPK activity, and positive regulation of smooth muscle cell proliferation (Figure 4D), consistent with our previous results (Figures 2M and 2N; Figures 3K and 3L). KEGG enrichment analysis demonstrated that the downregulated genes were involved in the MAPK signaling pathway, PI3K-Akt signaling pathway, and cytokine-cytokine receptor interaction (Figure S4G). Notably, the knockout of *Adma9* led to decreased expression levels of several metalloproteinase genes, suggesting that the function of extracellular matrix disassembly was suppressed in MOVAS cells incubated with rhCTHRC1 (Figure 4E). Meanwhile, the mRNA and protein levels of VSMC contractile markers were restored in *Adam9*-KO cells in the presence of rhCHTRC1 (Figures 4E and 4F; Figure S4H). Collectively, these findings provide evidence that CTHRC1 activates downstream dedifferentiation signals via ERK1/2 phosphorylation, dependent on its interaction with the ADAM9 on VSMCs.

Previous studies have shown that suppression of ADAM9 by a small molecule inhibitor (ADAM9i) restricted PDAC progression by blocking the KRAS/ERK signaling cascade.^71^ We therefore sought to evaluate the effect of ADAM9-specific inhibitor on CTHRC1-induced ERK1/2 signaling transduction in MOVAS cells. The ADAM9 inhibitor has a high binding affinity for ADAM9 and shows no inhibitory effects on other homologs within the ADAM family.^71^ After rhCTHRC1 stimulation, ADAM9 inhibitor-treated cells were collected for RNA-seq and immunoblotting. As anticipated, pharmacological inhibition of ADAM9 significantly rescued the VSMC contractile phenotype and simultaneously downregulated MMP gene expression by restraining downstream ERK pathway activation (Figures 4G-4I; Figures S4I-S4K). Furthermore, we confirmed the interaction between CTHRC1 and ADAM9 in human aortic smooth muscle cells (HASMCs) (Figure 4J). ADAM9i treatment indeed elevated contractile markers and inhibited downstream ERK signals in cultured HASMCs (Figure 4K). These data indicate that rewiring the CTHRC1-ADAM9-ERK1/2 axis can suppress the phenotypic switching of contractile SMCs, potentially restricting AD development.

### CTHRC1 blockade therapy protects against aortic dissection in mice

Given that CTHRC1 is an exocrine protein, its function could be blocked by specific antibodies. We used purified CTHRC1 protein as an immunizing antigen to generate anti-CTHRC1 monoclonal antibodies (anti-CTHRC1 Ab). The purified CTHRC1 protein showed high amino acid sequence homology with both human and mouse CTHRC1 proteins (Figure S5A). The sequences of the antibody heavy and light chains were confirmed through variable region cloning, expressing, and sequencing (Figure 5A). Additionally, molecular docking analysis showed 12 common amino acids in the CTHRC1 binding sites that interact with both the variable region of anti-CTHRC1 Ab and the extracellular structure of ADAM9 (Figure 5B). ELISA and immunoblotting assays indicated the high affinity, specificity, and efficacy of the anti-CTHRC1 Ab (Figure S5B). After confirming the quality of the antibody *in vitro*, we treated WT mice intraperitoneally with either anti-CTHRC1 Ab or IgG Isotype antibody to investigate their effects on AD formation (Figure 5C). No mortality was observed in the PBS groups treated with either anti-CTHRC1 Ab or IgG Isotype during the study period, indicating the safety of the antibodies (Figure 5D). Notably, mice treated with anti-CTHRC1 Ab showed a significantly higher survival rate compared to the IgG isotype group (Figure 5D), suggesting a strong therapeutic effect of anti-CTHRC1 Ab on AD. Ultrasonography also revealed a significant decrease in the maximal diameter of aortas from the anti-CTHRC1 Ab group (Figures 5E and 5F). Approximately 60% of mice treated with anti-CTHRC1 Ab developed AD, whereas 100% of mice in the IgG isotype group exhibited AD (*P*value ≤ 0.05) (Figures 5G and 5H). The microscopic structure of aortas from the anti-CTHRC1 Ab group displayed fewer pathological features, indicating its therapeutic effect on vascular remodeling (Figures 5I and J). These findings align with the results observed upon *Cthrc1* knockout (Figures 2C to 2I), supporting the potential for precise intervention in AD.

**Figure 5.**
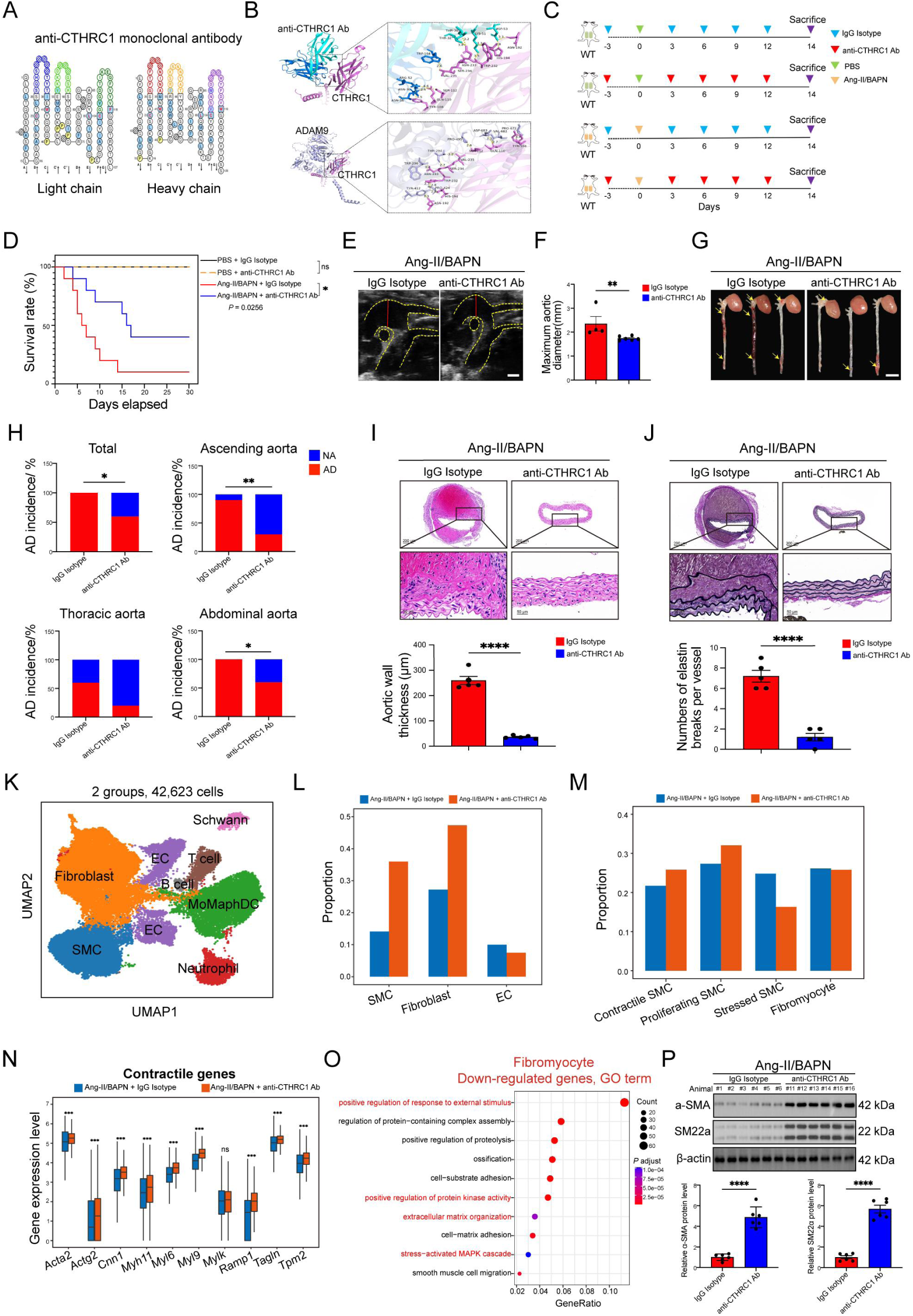
CTHRC1 blockade therapy protects against aortic dissection in mice. (A) The sequences of heavy and light chains from anti-CTHRC1 monoclonal antibody (anti-CTHRC1 Ab), confirmed by variable region cloning, expressing and sequencing. (B) Molecular docking analysis showing the interaction between antibody-CTHRC1 antigen and CTHRC1-ADAM9 receptor at the molecular level. (C) Experiment design. WT mice were intraperitoneally injected with anti-CTHRC1 Ab or IgG Isotype at a dose of 5 μg/g body weight under saline or Ang-II/BAPN administration (n=10 per group). (D) Kaplan-Meier survival rates analysis of the mice treated with anti-CTHRC1 Ab or IgG Isotype under Ang-II/BAPN administration. (Kaplan-Meier survival rates analysis, \**P* ≤ 0.05, ns, no significance.) (E) Representative ultrasound images of aorta in mice treated with anti-CTHRC1 Ab or IgG Isotype under Ang-II/BAPN administration (scale bar, 1mm). (F) Measurements of maximum aortic diameter of mice in E (n=6-9 per group). (Student’s t-test, \*\**P* ≤ 0.01.) (G) Representative pictures of whole aortas in mice treated with anti-CTHRC1 Ab or IgG Isotype under Ang-II/BAPN administration. The yellow arrows indicate the dissected sites of vascular lesions. (H) Quantification of AD incidence in the whole aorta, ascending, thoracic and abdominal aortas from the mice in E. (Mann-Whitney test, \*\**P* ≤ 0.01.) (I) Representative images of histological staining with hematoxylin and eosin in aortic sections from the mice in E. Bottom: Measurements of aortic wall thickness of aortas (scale bar, 50 μm; scale bar, 200 μm). (Student’s t-test, \*\*\*\**P* ≤ 0.00001.) (J) Representative images of Elastica van Gieson staining in aortic sections from the mice in E. Bottom: Numbers of elastin breaks per vessel. (scale bar, 50 μm; scale bar, 200 μm). (Student’s t-test, \*\*\*\**P* ≤ 0.00001.) (K) K to O: scRNA-seq of the aortic tissues from WT mice treated with anti-CTHRC1 Ab or IgG Isotype under Ang-II/BAPN administration (n=2 per group). UMAP plot of the 42,623 single cells, displaying the identification of 8 major clusters. (L) The proportion of major cell types including SMCs, ECs and FBs in WT mice treated with anti-CTHRC1 Ab or IgG Isotype under Ang-II/BAPN administration (M) Bar diagram showing the percentage of 4 SMC subclusters in aortic SMCs of the two groups of mice. (N) Boxplot illustrating the differential expression of contractile genes in aortic SMCs of the two groups of mice. (\*\*\**P ≤* 0.001, ns, no significance.) (O) Gene Ontology enrichment analysis of differentially down-regulated genes in fibromyocytes of the two groups of mice. (P) Immunoblotting analysis and quantification of α-SMA and SM22α expression in aortas from the WT mice treated with IgG Isotype or anti-CTHRC1 Ab under Ang-II/BAPN infusion (n=6-9). (Student’s t-test, \*\*\*\**P* ≤ 0.00001.) See also Figure S5.

Furthermore, we performed RNA-seq at both single-cell and bulk levels on aortic tissues from Ang-II/BAPN-induced mice treated with either anti-CTHRC1 Ab or IgG isotype. Using scRNA-seq data, we identified 8 major cell clusters (Figure 5K). Distribution analysis showed that CTHRC1 blockade significantly altered the general proportion of major cell types, particularly SMCs, fibroblasts, and ECs (Figure 5L). Interestingly, the cell distribution pattern and gene expression profile in mice treated with anti-CTHRC1 Ab resembled the SMC phenotypic modulation observed upon *Cthrc1* knockout (Figures 3D to 3H). Four SMC subclusters were identified in the aortas, with an increased percentage of contractile SMCs and elevated expression of contractile genes (Figures 5M and 5N). Although the proportion of fibromyocytes did not change after the CTHRC1 blockade therapy, down-regulated genes in this subtype of SMC were associated with the stressed-activated MAPK cascade (Figure 5O). At the protein level, CTHRC1 blockade prevented Ang-II/BAPN-induced downregulation of contractile markers (SM22α and α-SMA) (Figure 5P). Collectively, these findings strongly suggest that blocking CTHRC1 with antibodies inhibits SMC phenotypic modulation in AD.

Bulk RNA-seq data revealed that down-regulated genes in the anti-CTHRC1 Ab group were associated with positive regulation of the Erk1 and Erk2 cascade, extracellular matrix disassembly, and positive regulation of smooth muscle cell proliferation (Figures S5C and S5D). In particular, contraction-related genes, such as *Acta2*, *Myocd*, and *Myl4* were up-regulated in the aortas from the anti-CTHRC1 Ab group, whereas key genes associated with extracellular matrix disassembly (*Mmp3*, *Mmp9*, *Mmp12*, and *Mmp13*) were down-regulated (Figures S5E). The therapeutic effects of anti-CTHRC1 Ab were also evaluated in female mice, and results showed comparable effects to those observed in male mice (Figures S5F to S5K). Therefore, our data from CTHRC1 blockade therapy and *Cthrc1* knockout confirmed the pivotal role of CTHRC1 in AD progression, highlighting its potential as a promising therapeutic target for AD.

### Serum CTHRC1 serves as a promising diagnostic biomarker for AD

After assessing the impact of CTHRC1 on AD pathogenesis and development using mouse models, we sought to explore whether CTHRC1 also plays an essential role in humans. Hence, we collected aortas from patients for scRNA-seq (Figure 6A). After quality control, we obtained 45,501 single cells in total from normal arteries (healthy controls) and AD aortas, identifying 7 major cell clusters (Figures S6A to S6C). In the fibroblasts of the dissected aortas, we observed a significantly elevated expression level of CTHRC1 (Figure 6B). We then examined SMC phenotypic alterations in aortic tissues between AD patients and healthy controls. The percentage of SMCs was significantly lower in human dissected aortas compared to normal arteries (Figure 6C), and the four SMC subpopulations identified in mouse models were also observed in human patients (Figure S6D). Further analysis of differentially expressed genes among the four SMC subclusters revealed that the expression levels of contractile genes were down-regulated in dissected aortas (Figure 6D and S6E). Consistently, the down-regulated genes in contractile SMCs were enriched in contraction function (Figure S6F). In fibromyocytes, GO analysis revealed that up-regulated genes were involved in the ERK1 and ERK2 cascade and positive regulation of smooth muscle cell proliferation in human AD (Figure 6E). Moreover, bulk RNA-seq data showed increased expression levels of extracellular matrix disassembly-related genes and decreased expressions of genes associated with SMC contraction in AD patients (Figure 6F and Figure S6G). Immunoblotting analysis demonstrated dramatically increased protein levels of CTHRC1, ADAM9, and phospho-ERK1/2 in AD tissues compared to controls (Figures 6G and 6H), indicating the activation of the CTHRC1-ADAM9-ERK signaling axis. Furthermore, immunofluorescence analysis confirmed lower levels of SM22α and α-SMA in human dissected aortas (Figure 6I). Taken together, these data suggest a conserved pathogenic mechanism involving CTHRC1 in both mice and humans.

**Figure 6.**
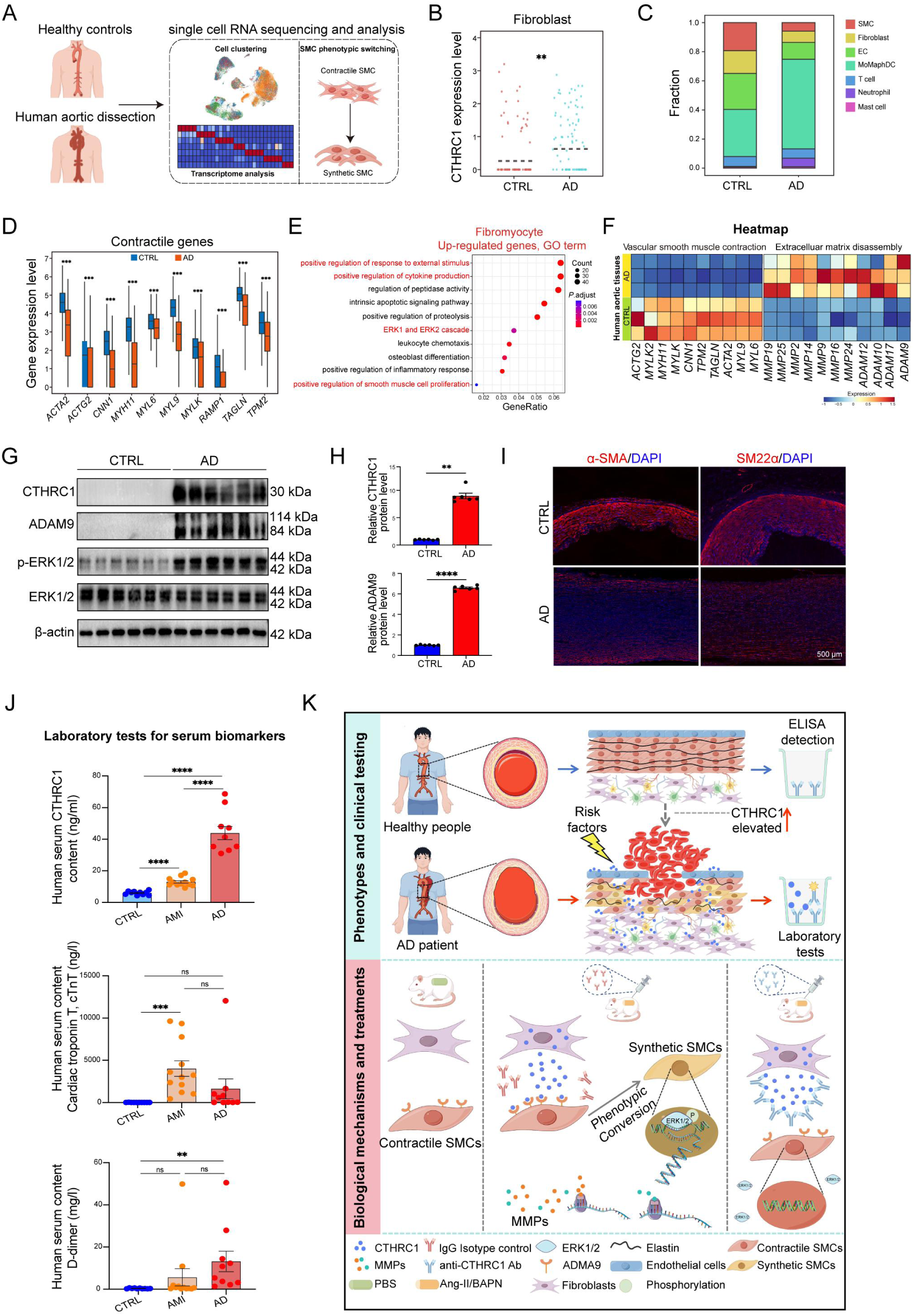
Serum CTHRC1 serves as a promising diagnostic biomarker for AD. (A) A to E: scRNA-seq of aortic tissues in healthy controls and human aortic dissection (n=2 per group). Experiment design of scRNA-seq. (B) Dotplot showing the expression level of CTHRC1 in fibroblasts of the aortas from AD patients and controls. Dash line in each group represents its mean expression value. (C) The proportion of 7 major cell clusters in AD patients and controls. (D) Boxplot displaying the differential gene expression of contractile genes in aortic SMCs of the controls and AD patients. (E) Gene Ontology enrichment analysis of differentially up-regulated genes in fibromyocytes from AD patients versus controls. (F) Heatmap showing the expression levels of genes related to vascular smooth muscle cell contraction and extracellular matrix disassembly in aortic tissues from AD patients and controls by bulk RNA-seq. (\*\*\**P ≤* 0.001.) (G) Immunoblotting analysis of CTHRC1, ADAM9, p-ERK1/2 expression levels in human aortas (n=6). (H) Quantification of CTHRC1, ADAM9, p-ERK1/2 protein expression levels in G. (Student’s t-test, \*\**P* ≤ 0.01, \*\*\*\**P ≤* 0.0001.) (I) Immunofluorescence staining of α-SMA and SM22α expression in human aortas (scale bar, 500 μm). (J) Laboratory tests for human serum CTHRC1, cTnT, and D-dimer contents in control individuals, acute myocardial infraction (AMI) and AD. (Student’s t-test, \*\**P* ≤ 0.01, \*\*\**P ≤* 0.001, \*\*\*\**P ≤* 0.0001, ns, no significance.) (K) Graphical abstract illustrating an important role of CTHRC1 acts as a candidate circulating biomarker and therapeutic target for aortic dissection through CTRHC1-ADAM9-ERK axis. See also Figure S6.

To determine whether CTHRC1 serves as a biochemical marker for diagnosing AD, we measured CTHRC1 levels in human serum. We found that CTHRC1 was significantly elevated in the peripheral blood of patients with AD (Figure 6J). According to published data, CTHRC1 produced by cardiac fibroblasts responds to cardiac injury and participates in myocardial remodeling after acute myocardial infarction (AMI).^72^ Thus, we assessed the serum level of CTHRC1 alongside other commonly used clinical testing markers, including cTnT, D-dimer, myoglobin, BNP, hCRP, and CKMB, in both AMI and AD patients (Figures 6J; S6H and S6I). CTHRC1 levels were approximately 2.1-fold higher in AMI patients than in controls, consistent with previous studies.^72^ Notably, CTHRC1 levels in AD patients were much higher than those in AMI patients (with an increase of about 3.3 fold) (Figure 6J). Unlike existing clinical biomarkers such as D-Dimer and cTnT (Figure 6J), serum CTHRC1 levels can accurately and straightforwardly distinguish between AD and AMI patients, making it a promising novel biomarker for the differential diagnosis of patients with emergency chest pain. Based on these intriguing animal and clinical data, we developed a human CTHRC1 ELISA kit for clinical diagnosis in future study (Figure S6J). In summary, these findings highlight that the altered expression level of CTHRC1 positions it as a potential novel blood biomarker for the diagnosis of AD, providing a non-invasive and easy-to-use diagnostic tool with significant clinical implications.

## Discussion

Aortic dissection (AD) remains a significant challenge in emergency diagnosis. Despite its rare occurrence, its lethality is high, and delays in prompt and accurate diagnosis and treatment often results in fatal outcomes from aortic rupture or cardiac tamponade.^73,74^ Previous studies indicate that there have been minimal advancements in accurate diagnosis and timely intervention.^75^ Herein, we challenged both male and female mouse models with Ang-II combined with BAPN via mini-osmotic pumps to replicate the human condition. Notably, gene expression patterns in these mouse models closely parallel those observed in human AD. Utilizing this reliable model, we have demonstrated that adventitial fibroblasts exocrine high levels of CTHRC1 into peripheral blood during AD development, suggesting its potential as a blood-based biomarker. Moreover, targeted therapy by anti-CTHRC1 antibodies ameliorated AD formation by inhibiting the transition of VSMCs from a contractile to a synthetic phenotype, a well-defined VSMC population important for vascular remodeling.^20^ At the molecular level, secreted CTHRC1 interacts with ADAM9 on VSMCs, leading to the activation of the ERK1/2 signaling pathway and downregulation of contractile gene expression. Our findings elucidates the role of CTHRC1 in modulating intercellular communication between fibroblasts and VSMCs through the CTHRC1-ADAM9-ERK1/2 signaling axis, thereby aiding in the identification of AD patients among individuals presenting with acute chest pain.

Over the past decades, CTHRC1 has emerged as a key biomarker and contributor to fibroblast associated pathological changes, including myocardial remodeling,^72^ tumor immunosuppression,^76^ and organ fibrosis.^77,78^ CTHRC1 is a secreted 28-kDa glycoprotein, highly conserved in mammals^79^ and predominantly expressed in fibroblasts and fibroblast-like cells.^80,81^ Previous studies have shown that *Cthrc1^+^* fibroblasts express high level of collagens, which are essential for homeostasis and pathological fibrosis.^77,79,82^ This finding aligns with the phenotype observed in AD mouse models with *Cthrc1* inhibition. Simultaneously, analysis of public datasets based on patient aortic aneurysm tissues^47^ reveals that rupture-prone aortic aneurysms exhibit remarkably elevated level of *CTHRC1* (Figure 1J). Notably, our data indicated that both *Cthrc1* deletion and antibody treatment significantly suppressed AD formation and prolonged survival in mice, regardless of gender. This suggests that the effects of CTHRC1 on AD may be mediated through the reprogramming and remodeling of the vascular microenvironment, positioning CTHRC1 as a potential therapeutic target. Additionally, our study revealed that targeted therapy against CTHRC1 could suppress AD by impeding fibroblast-mediated phenotypic transformation of VSMCs through the CTHRC1-ADAM9-ERK axis. However, MMPs, cytokines, and chemokines regulated by this pathway may affect not only VSMCs but also immune and endothelial cells within the vascular microenvironment. For instance, MMP9 has been identified as an activator of SMC migration and proliferation,^83,84^ but it also influences macrophage polarization,^85^ neutrophil migration,^86^ and endothelial cell migration^87^ under certain conditions. Therefore, it is crucial to investigate the effects of MMPs or cytokines regulated by CTHRC1-ADAM9 interactions on other cell types. Notably, CTHRC1 is expressed not only in pathological fibrosis but also during the healing process following injury, such as myocardial infarction.^72^ Future studies should evaluate the systemic effects of CTHRC1 inhibition in vivo and in humans, particularly when CTHRC1 antibodies or inhibitors are used.

Our findings demonstrate that ADAM9 serves as the binding protein for CTHRC1 on VSMCs, mediating AD formation and progression through the regulation of ERK1/2 signaling. The ADAM family of proteins is involved in various biological processes, including cell fate determination, wound healing, heart development, cell migration, sperm-egg interaction, immunity, cell proliferation, and angiogenesis.^88,89^ Recently, the role of ADAMs and ADAMTSs in the development of AD has garnered increasing attention, with several members of these families (e.g. ADAM10, ADAM15, ADAMTS1, ADAMTS16, ADAMTS18),^50,51,90,91^ exhibiting differential expressions in case of AD or aneurysm.^50^ In this study, we demonstrated the effects of CTHRC1 on VSMC transformation through the ADAM9-mediated ERK1/2 signaling cascade in AD. It is noteworthy that ADAMs and ADAMTs family members, such as ADAM8, ADAM15, ADAM17, ADAMTS1, and ADAMTS5 have been widely reported to be engaged in ERK1/2 signaling.^66-69,92^ Additionally, genetic ablation or pharmacological inhibition of ADAM9 disrupts the functions of CTHRC1, leading to increased expression of SMC contractile genes and decreased expression of extracellualr matrix genes. Correspondingly, previous studies have shown elevated ADAM9 expression in SMCs associated with aortic aneurysm.^70^ While ADAM9 was the most prominent ADAM family member identified in our study through IP-MS analysis, it is not the only ADAM of interest. For example, transcriptomic data revealed significantly increased expressions of ADAM8 and ADAM17 in mouse and human AD samples, alongside decreased *Adam12* expression in *Cthrc1^-/-^* mice. Hence, future studies should systematically evaluate the role of ADAMs in the development of AD.

Clinically, especially in emergency room (ER), distinguishing between chest pain due to acute myocardial infarction (AMI) and that related to AD presents a significant challenge for emergency physicians. AMI and AD can share similar symptoms, such as squeezing, tightness, stabbing, or dull pain, as well as signs like elevated heart rate and blood pressure, and even laboratory results, including elevated troponin, D-dimer, C-reactive protein. Notably, patients with AMI require anti-coagulation therapy, which poses a risk of fatal bleeding in AD cases due to the high likelihood of aortic rupture. In some instances, proximal AD may affect coronary arteries, potentially mimicking AMI. While contrast-enhanced computed tomography (CT) is highly accurate for diagnosing AD with notable sensitivity and specificity,^93^ it remains time-consuming and relatively low cost-effective, with only 3 out of every 1,000 patients presenting with chest pain utimately diagnosed with AD.^94^ Therefore, a straightforward and specific blood test is urgently needed to expedite the diagnosis of AD and ensure appropriate clinical management. Our clinical data indicate that CTHRC1 is a sensitive and easily measurable blood biomarker that can differentiate between AD and AMI. It is noteworthy that blood CTHRC1 concentrations were also significantly higher in patients with rheumatoid arthritis (RA) compared to healthy individuals.^95^ However, in contrast to RA, AD is an emergency that presents with completely distinct symptoms or signs. Next, we plan to enroll a larger cohort of patients in a multi-center prospective study to establish an unbiased optimal cut-off point for CTHRC1 using clinical kits and determine the peak levels of CTHRC1 in AD. Additioanlly, developing a point-of-care testing (POCT) device for CTHRC1 that provides rapid, reliable results from a single drop of blood in under 10 minutes in the ER will be essential for managing AD (Figure S6K), similar to existing tests for cTnT in AMI or D-dimer in thrombosis. This wold facilitate the assessment of chest pain during ambulance transport or in the emergency room, allowing for timely and appropriate patient care (Figure S6L).

Our observation demonstrate that inhibition of CTHRC1 secreted by vascular fibroblasts diminishes the remarkable phenotypic plasticity of SMCs, the formation of SMC-associated MMPs and cytokines, reduces vascular remodeling, and maintains arterial homeostasis. We found that the effects of CTHRC1 on gene regulation are largely mediated by the membrane-anchored protein ADAM9 on VSMCs. However, the mechanisms underlying ADAM9’s regulation of ERK1/2 and its downstream gene expression require further investigation. Our preliminary data suggest that *ADAM9* gene expression can be regulated in *vivo* and *in vitro* at intensity in the presence of CTHRC1 protein, suggesting that CTHRC1 may influence ADAM9 through transcriptional regulation and protein-protein interactions. Further studies are needed to gain insight into these regulatory mechanisms. Moreover, we developed a highly specific antibody that effectively blocks CTHRC1, underscoring its immediate translational potential in AD treatment. However, further research is necessary to determine whether anti-CTHRC1 antibodies can also suppress the progression of aortic aneurysms or recurrent aortic dissection (RAD). We anticipate that CTHRC1-targeted therapies against can be developed in the near future, enhancing treatment options for AD, RAD, and aortic aneurysm.

## ACKNOWLEDGMENTS

This study was supported by the National Natural Science Foundation of China (No. 82372783, 32170660, 82171812, 82273018, 82272685, 81900306 and 82472745), the National Key Research and Development Program of China (No. 2023YFC3405200 and 2024NSFSC0734), the Natural Science Foundation of Sichuan Province (No. 2023NSFSC0580 and 2023YFS0128), the Shanghai Outstanding Academic Leaders Plan, the Sichuan Province Tianfu Emei Project, the Natural Science Foundation of Shanghai (No. ZR1492800), Shanghai Pilot Program for Basic Research, the Fundamental Research Funds for the Central Universities (No. 22120240435). We would like to thank Dr Zhigang Zhang and Dr Jun Li from Shanghai Jiaotong University for sharing their antibody. We also thank Dr Shirley X. Liu and Dr. Tengfei Xiao from Shanghai Xunbaihui Biotechnology for their valuable discussions, and we thank Dr Hailing Cheng from Dalian Medical School for her help with editing and preparing this manuscript. Cartoons in graphical abstract were created with figdraw.com.

## AUTHOR CONTRIBUTIONS

X.W., and C.W., conceived, designed, and initiated the study. J.C., J.L., and P.W., performed most experiments. L.J., analyzed RNA-seq and scRNA-seq data. H.H., C.C., and X.D., analyzed proteomics data. Q.L., T.Y., J.T., J.L., X.W., Y.Z., L.L., H.H., C.L., L.W., Y.X., Z.C., J.Z., T.L., Y.C., P.L., Y.C., J.F., J.L., L.W., and L.L., enrolled the human subjects of the study. X.Z., G.X., Z.D., C.S., and C.X., provided suggestions for experiments in vivo. G.X., C.J., and H.W., provided assisteneces with production of antibodies. J.Z., and Z.W., provided suggestions for single-cell RNA-seq data analyses. X.W., W.T., and C.W. jointly prepared the manuscript with input from all authors. X.W., W.T., and C.W., secured funding and supervised the work.

---

**Figure S1.**
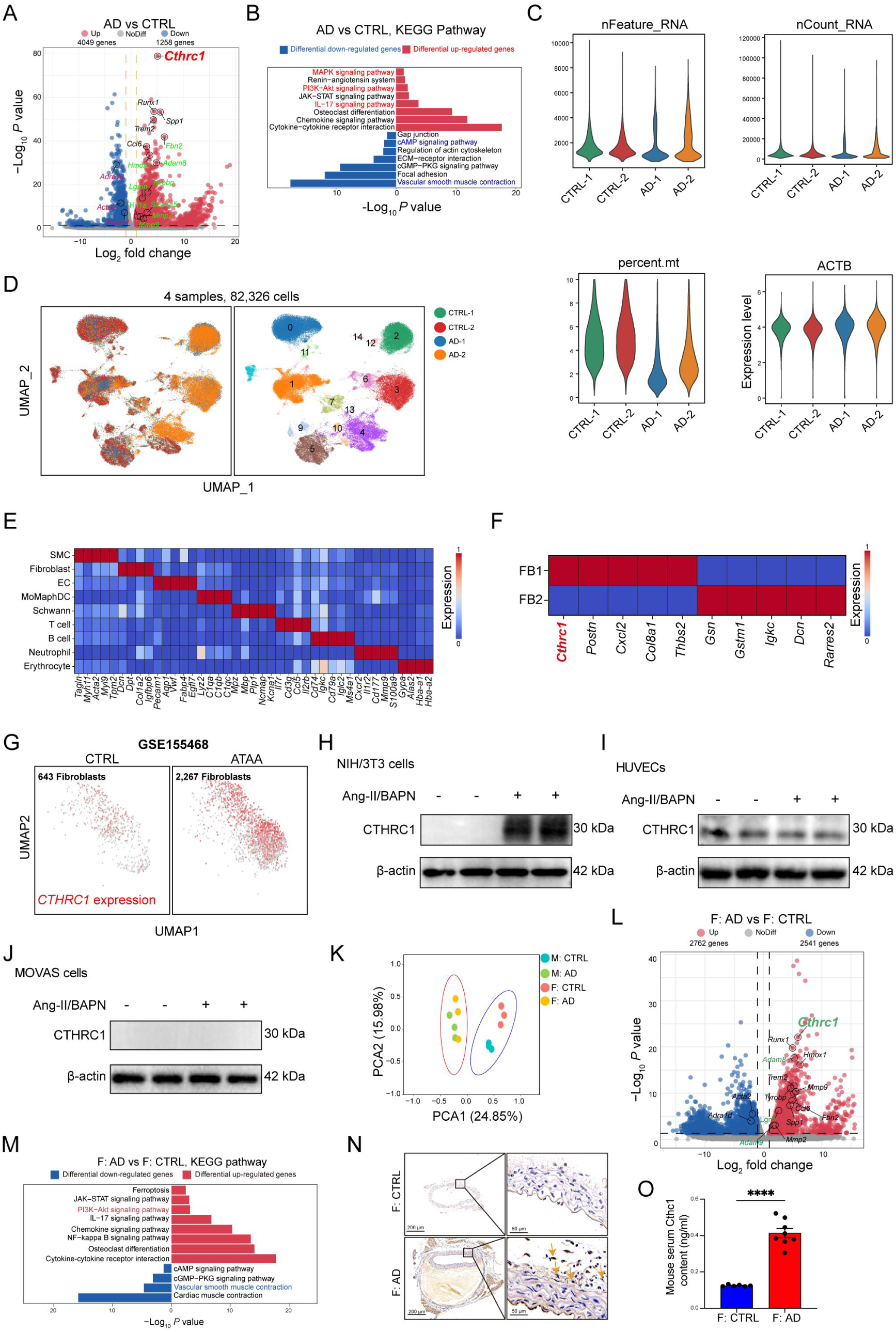
CTHRC1 is highly expressed in the aortas of mouse models and patients with AD, related to Figure 1. (A) A and B: bulk RNA-seq of aortic tissues from control and AD mice (n=3 per group). Volcano plot displaying differentially expressed genes in aortas between the two groups. *Cthrc1* was significantly up-regulated in AD mouse models. (B) KEGG enrichment analysis of differentially expressed genes in aortic tissues from control and AD mice. (C) C to F: scRNA-seq data of 4 samples from control and AD mice. Violin plots showing nFeature, nCount, percent.mt and expression level of the housekeeping gene beta-actin (Actb) after quality control of 4 samples. (D) UMAP plot of 82,326 single cells from 4 samples colored according to the 15 clusters. (E) Heatmap showing expression signatures of the marker genes in each cell type. (F) Heatmap showing expression signatures of the top differentially expressed genes in 2 fibroblast subclusters. (G) UMAP plots showing *CTHRC1* expression in fibroblasts from control individuals and patients with ascending thoracic aortic aneurysm (GSE155468). (H) H to J: CTHRC1 protein levels in vitro (NIH3T3 cells (H), human umbilical vein endothelial cells (HUVECs) (I), and MOVAS cells(J), incubated with Ang-II (10^-6 M) and BAPN (0.2 mM). (K) bulk RNA sequencing of aortas in male and female mice. PCA of gene profiles of transcriptome between different sexes. (L) Volcano plot displaying differentially expressed genes in aortic tissues from control and AD female mice (n=3 per group). *Cthrc1* was also significantly up-regulated in AD female mouse models. (M) KEGG enrichment analysis of the differentially expressed genes in AD female mouse models. (N) Immunohistochemical staining for CTHRC1 in female mouse aortas. The orange arrows indicate CTHRC1 expression in adventitial fibroblasts of the dissected aortas. (O) Female mouse serum CTHRC1 content in control and AD mice (n=6). (an unpaired two-tailed Student’s t-test, \*\*\*\**P ≤* 0.0001.)

**Figure S2.**
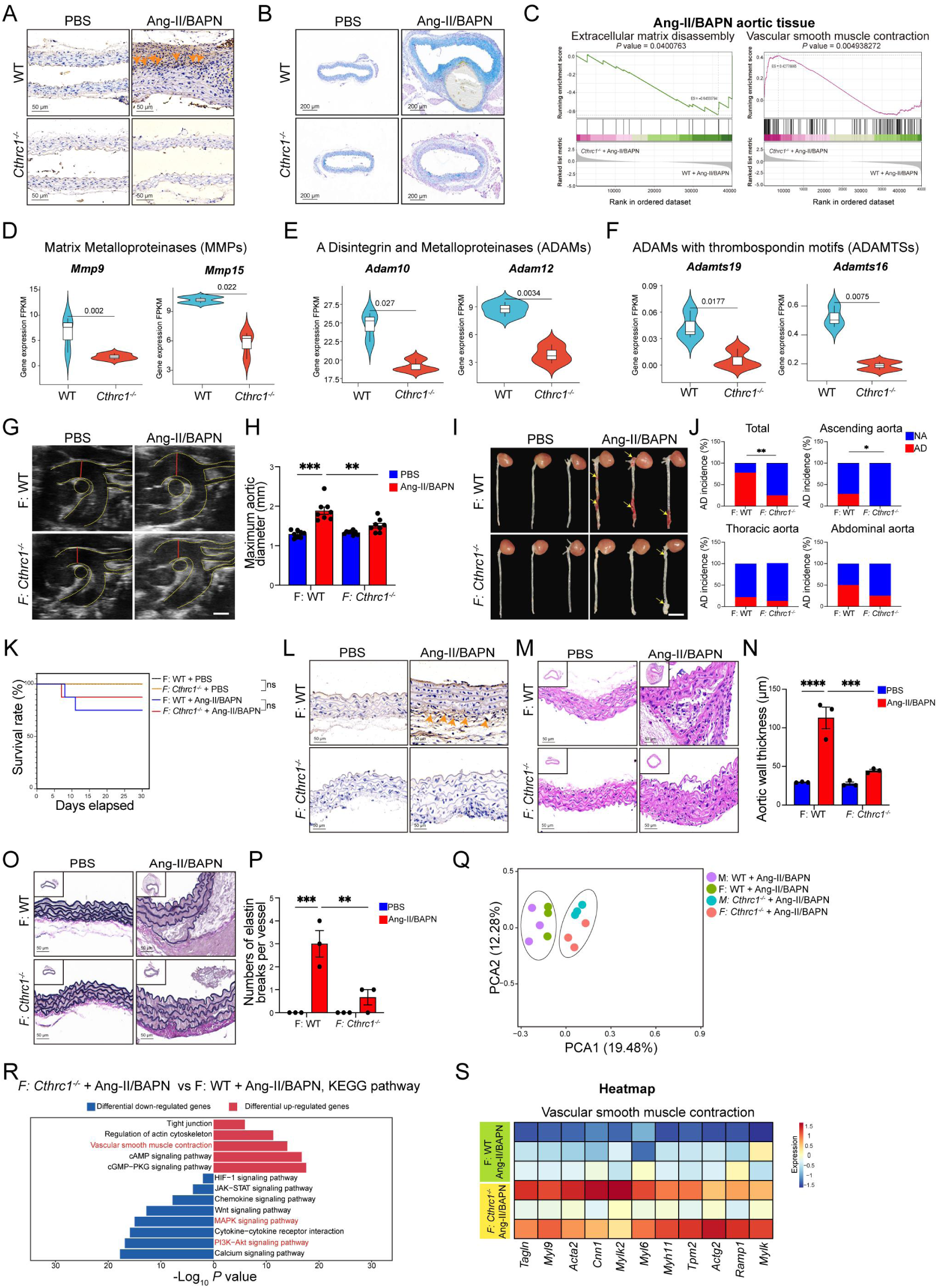
*Cthrc1* KO attenuates AD formation in mouse models, related to Figure 2. (A) Immunohistochemical staining for CTHRC1 in aortas from the *Cthrc1^-/-^* and WT mice. The orange arrows indicate CTHRC1 expression in adventitial fibroblasts of the dissected aortas, with no positive staining observed in *Cthrc1^-/-^* group. (B) Alican blue staining of aortas from the *Cthrc1^-/-^* and WT mice. (C) C to F: bulk RNA-seq of aortic tissues from WT and *Cthrc1^-/-^* mice. GSEA showing the inhibition of extracellular matrix disassembly and the activation of vascular smooth muscle contraction. (D) : D to F: Violin plots illustrating gene expressions of MMPs (D), ADAMs (E) and ADAMTSs (F) between WT and *Cthrc1^-/-^* mice infused with Ang-II/BAPN. MMPs: matrix metalloproteinases; ADAMs: a disintegrin and metalloproteinases; ADAMTSs: ADAMs with a thrombospondin motif. (G) G to P: 6-week-old female *Cthrc1^-/-^* mice (n=16) and WT mice (n=18) were treated with saline or Ang-II/BAPN for 14 days. Representative ultrasound images of aorta in female mice (scale bar, 1mm). (H) Measurements of maximum aortic diameter of mice in G (n=8 per group). (a Welch ANOVA test followed by a post hoc analysis using the Tamhane T2 method, \*\*\**P* ≤ 0.001; \*\**P* ≤ 0.01.) (I) Representative pictures of whole aortas from the female mice. The yellow arrows indicate the dissected sites of vascular lesions. (J) Quantification of AD incidence in the whole aorta, ascending, thoracic and abdominal aortas from WT and *Cthrc1^-/-^* female mice. (Fisher’s exact test, \**P* ≤ 0.05.) (K) Kaplan-Meier survival rates analysis of the four groups of female mice. (Kaplan-Meier survival rates analysis, ns, no significance.) (L) Immunohistochemical staining for CTHRC1 in aortas from the female mice. The orange arrows indicate CTHRC1 expression in adventitial fibroblasts of the dissected aortas from the WT female mice, with no positive staining observed in Cthrc1^-/-^ female mice (scale bar, 50 μm). (M) Representative images of histological staining with hematoxylin and eosin in aortic sections from the female mice (scale bar, 50 μm). (N) Measurements of aortic wall thickness of aortas. (a ordinary one-way ANOVA followed by Bonferroni’s multiple comparison test, \*\*\**P* ≤ 0.001; \*\*\*\**P* ≤ 0.0001.) (O) Representative images of EVG staining in aortic sections from the female mice (scale bar, 50 μm). (P) Measurements of numbers of elastin breaks per vessel. (a ordinary one-way ANOVA followed by Bonferroni’s multiple comparison test, \*\**P* ≤ 0.01; \*\*\**P* ≤ 0.001.) (Q) Q to S: bulk RNA-seq of aortic tissues from WT and *Cthrc1^-/-^* female mice. PCA of gene profiles of transcriptome between different sexes. (R) KEGG enrichment analysis of differentially expressed genes in aortic tissues from WT and *Cthrc1^-/-^* female mice. (S) Heatmap showing the expression levels of genes related to vascular smooth muscle cell contraction.

**Figure S3.**
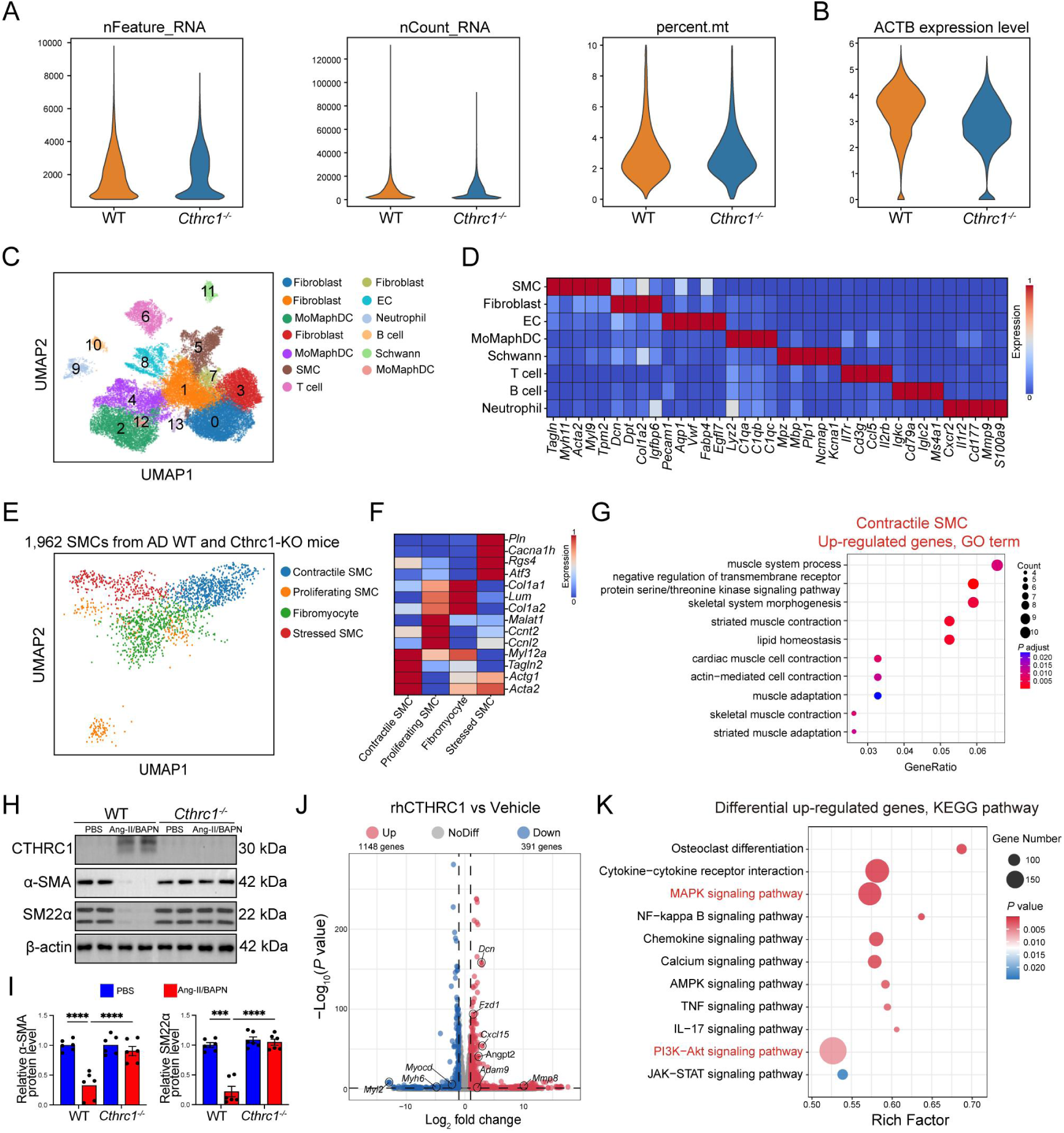
Systematic transcriptome analysis reveals communication between fibroblasts and VSMCs related to Figure 3. (A) A to G: scRNA-seq data of aortas from WT and *Cthrc1^-/-^* mice challenged with Ang-II/BAPN for 14 days. Violin plots showing data characteristics on nFeature, nCount and percent.mt after quality control of the two groups. (B) The violin plot showing the expression level of the housekeeping gene beta-actin (Actb) between the two groups. The relative stability of Actb expression indicates consistency. (C) UMAP plot of 82,326 single cells from the samples colored according to the 14 clusters. (D) Heatmap showing expression signatures of the marker genes in each cell type. (E) UMAP plot of 1,962 SMCs colored according to the identified 4 SMC subclusters. (F) Heatmap showing the scaled mean expression of signature genes in 4 SMC subclusters. (G) Gene Ontology enrichment analysis of up-regulated genes in contractile SMCs of the two groups of mice. (H) Immunoblotting analysis of α-SMA and SM22α expression in aortas from Ang-II/BAPN-infused WT and *Cthrc1^-/-^* mice (n=6). (I) Quantification of α-SMA and SM22α protein expression levels. (a ordinary one-way ANOVA followed by Bonferroni’s multiple comparison test, \*\*\**P* ≤ 0.001, \*\*\*\**P* ≤ 0.0001.) (J) J and K: whole transcriptome analysis of MOVAS cells treated with vehicle and rhCTHRC1. A volcano plot displaying differentially expressed genes. Upward expression of *Dcn*, *Adam9*, *Angpt2* and *Mmp8* gene in the presence of rhCTHRC1. (K) KEGG enrichment analysis of up-regulated genes in aortas from Ang-II/BAPN-infused WT and *Cthrc1^-/-^* mice.

**Figure S4.**
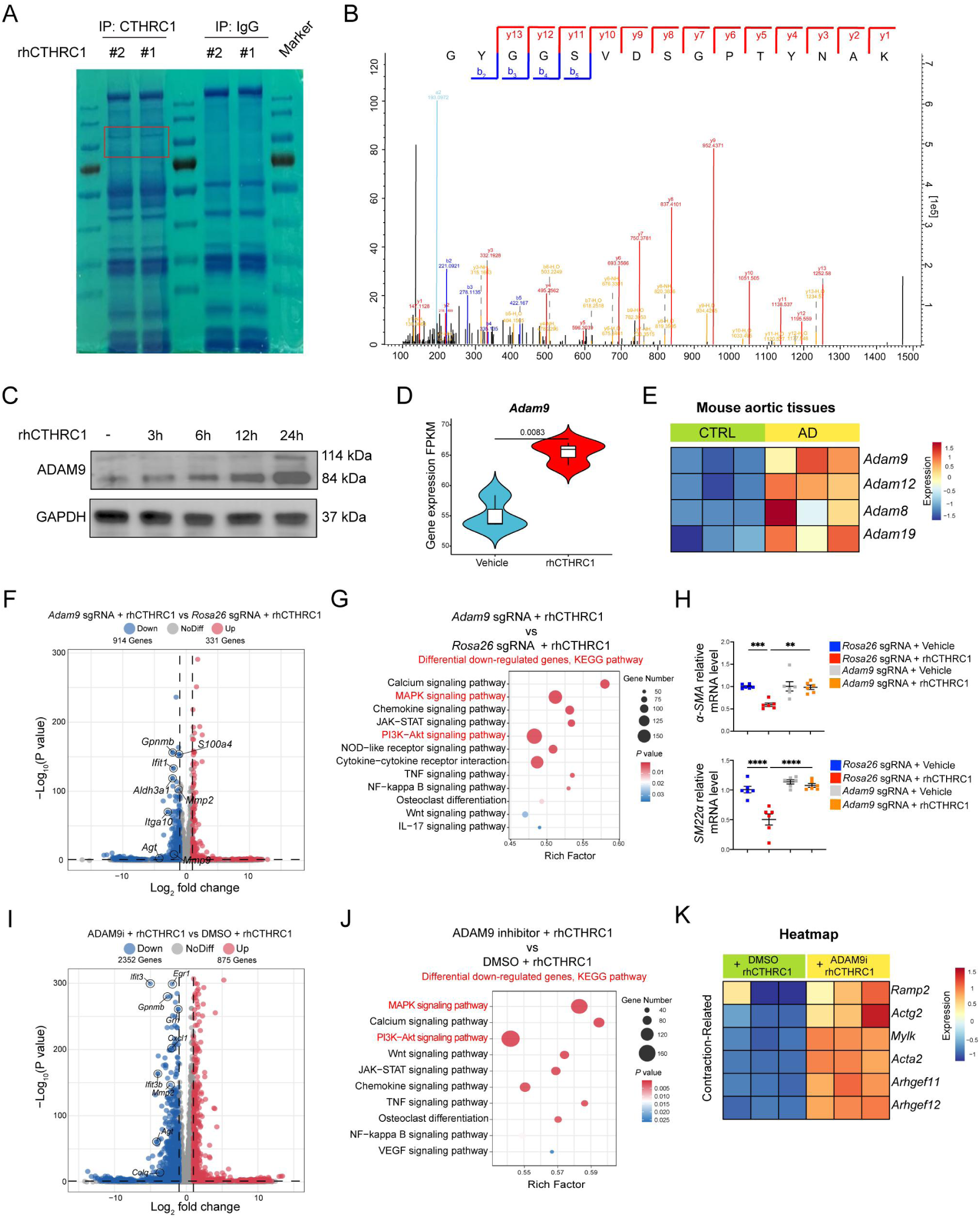
CTHRC1 regulates phenotypic switching of VSMCs by binding with ADMA9, related to Figure 4. (A) Coomassie staining revealing the protein content in the samples coimmunoprecipitated with CTHRC1. Red box indicates the potential interactors. (B) The representative peptide mass spectrum of ADAM9 receptor on VSMCs. (C) Immunoblotting analysis of ADAM9 in MOVAS incubated with rhCTHRC1 for 0, 3, 6, 12, and 24 hours. (D) Violin plot of *Adam9* gene expression in MOVAS cells with treatment of vehicle or rhCTHRC1. (E) Heatmap depicting the gene expression of ADAMs in control and AD mice. (F) F and G: transcriptome analysis of gene expression profiles in MOVAS cells without or with rhCTHRC1, knockouting *Roas26* or *Adam9*. A volcano plot of differentially expressed genes. The expression levels of *Agt*, *Mmp2* and *Mmp9* are down-regulated. (G) KEGG enrichment analysis of down-regulated genes in MOVAS cells in the presence of rhCTHRC1, knockouting *Roas26* or *Adam9*. (H) Relative mRNA levels of *α-SMA* and *SM22α* in MOVAS cells, without or with rhCTHRC1, knockouting *Roas26* or *Adam9*. (a ordinary one-way ANOVA followed by Bonferroni’s multiple comparison test, \*\**P ≤* 0.01, \*\*\**P* ≤ 0.001, \*\*\*\**P* ≤ 0.0001.) (I) I to K: Comprehensive profiles of genes in MOVAS cells without or with treatment of ADAM9i by bulk RNA-seq. A volcano plot of differentially expressed genes. *Egr1*, *Mmp2* and *Agt* are marked with down-regulated expression. (J) KEGG enrichment analysis of down-regulated genes in MOVAS cells in the presence of rhCTHRC1, with or without ADAM9i. (K) Heatmap showing the expression levels of contraction-related genes in MOVAS cells without or with treatment of ADAM9i, incubated with rhCTHRC1.

**Figure S5.**
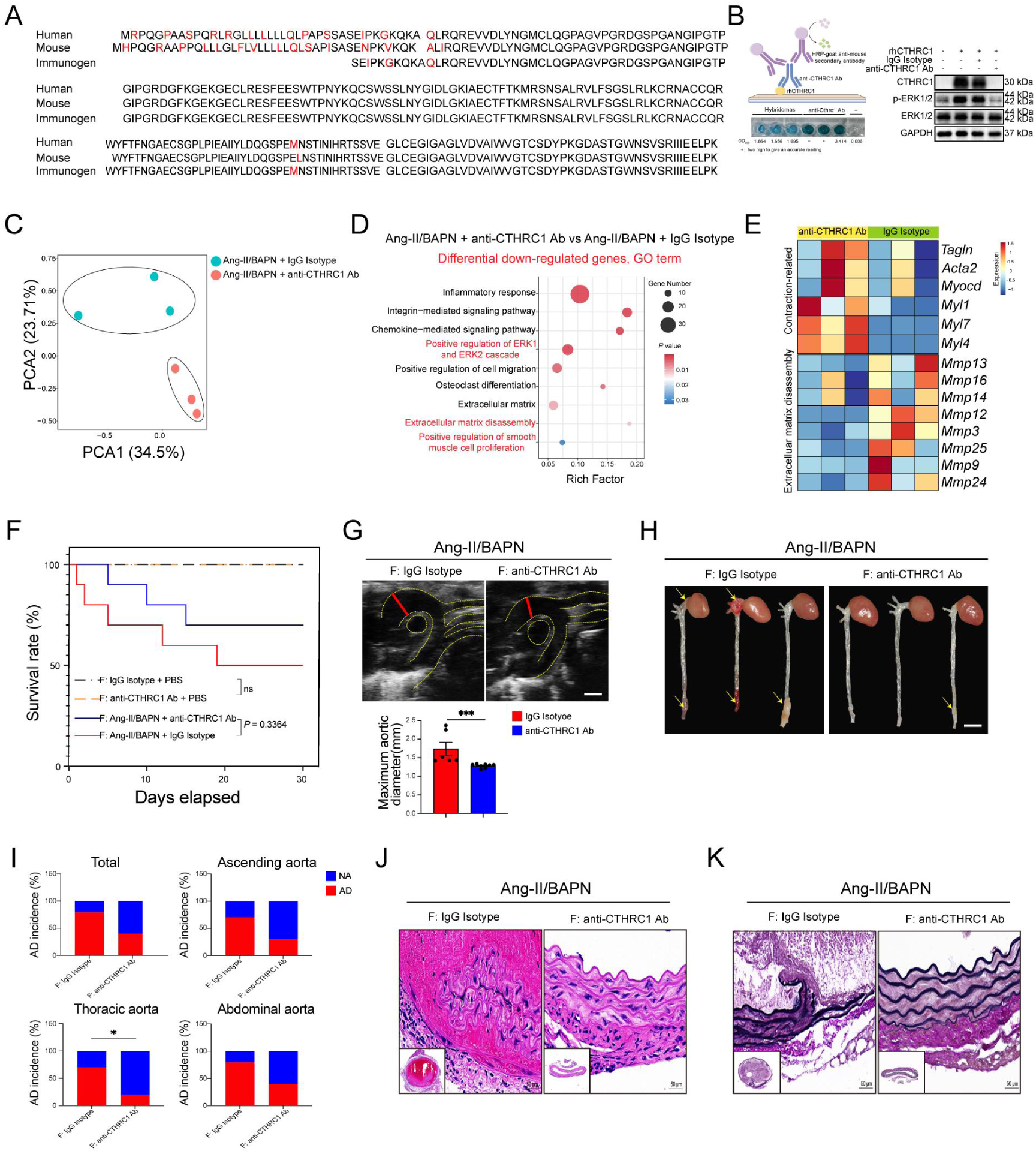
CTHRC1 blockade therapy protects against aortic dissection in mice, related to Figure 5. (A) Amino acid sequence homology of immunogen with human or mouse CTHRC1 protein. (B) The affinity, specificity, and efficacy of anti-CTHRC1 Ab confirmed by ELISA (left) and immunoblotting (right) analysis. (C) C to E: mRNA sequencing results illustrating the transcriptomic profiles of aortas from WT mice treated with IgG Isotype and anti-CTHRC1 Ab (n=3 per group). PCA of transcriptome profiles in aortic tissues between the two groups. (D) Gene Ontology enrichment analysis of down-regulated genes between the two groups (anti-CTHRC1 Ab versus IgG Isotype). (E) Heatmap showing the expression levels of genes related to vascular smooth muscle cell contraction and extracellular matrix disassembly. (F) F to K: WT female mice were intraperitoneally injected with anti-CTHRC1 Ab or IgG Isotype under saline or Ang-II/BAPN administration (n=10 per group). Kaplan-Meier survival rates analysis of the four groups of female mice. (Kaplan-Meier survival rates analysis, ns, no significance.) (G) Upper: Representative ultrasound images of aorta in female mice (scale bar, 1mm). Bottom: Measurements of maximum aortic diameter of mice (n=6 per group). (Mann Whitney test, \*\*\**P* ≤ 0.001.) (H) Representative pictures of whole aortas in female mice treated with IgG Isotype or anti-CTHRC1 Ab. (I) Quantification of AD incidence in the whole aorta, ascending, thoracic and abdominal aortas in female mice. (Fisher’s exact test, \**P* ≤ 0.05.) (J) Representative images of histological staining with hematoxylin and eosin in aortic sections from the female mice. (K) Representative images of EVG staining from the female mice injected with anti-CTHRC1 Ab or IgG Isotype.

**Figure S6.**
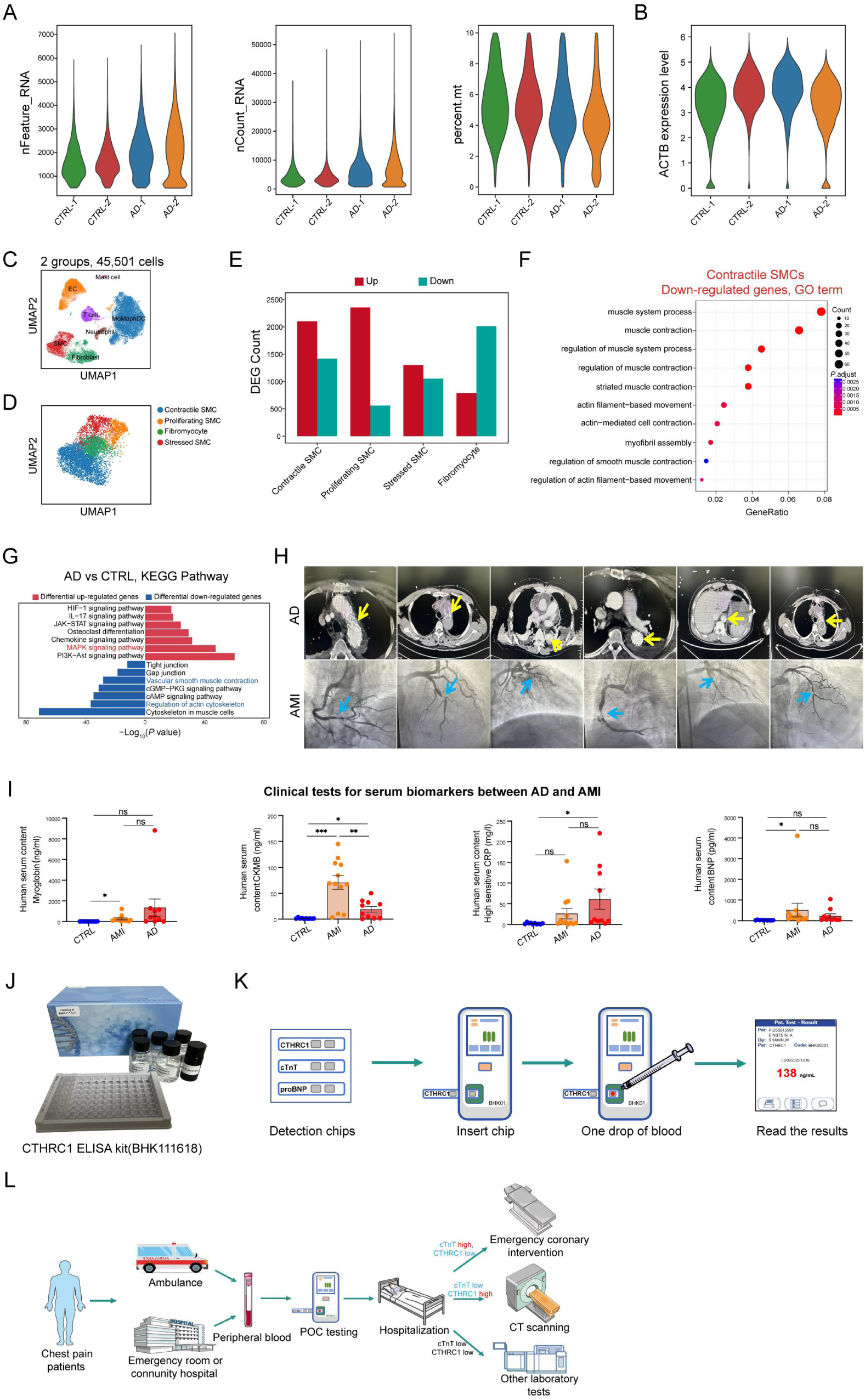
Serum CTHRC1 serves as a promising diagnostic biomarker for AD, related to Figure 6. (A) A to E: scRNA-seq data of aortas from human healthy controls and AD patients (n=2 per group). Violin plots showing data characteristics on nFeature, nCount and percent.mt after quality control of the two groups. (B) Violin plot showing the expression level of the housekeeping gene beta-actin (ACTB) of the four samples. The relative stability of ACTB expression indicates consistency. (C) UMAP plot of 45,501 single cells from 4 samples colored according to 7 different cell clusters. (D) UMAP plot of SMCs colored according to the identified 4 SMC subclusters. (E) Bar graphs depicting the numbers of differentially expressed genes in SMC subclusters in healthy controls and AD patients. (F) Gene Ontology enrichment analysis of down-regulated genes in contractile SMCs of the aortas. (G) KEGG enrichment analysis of differentially expressed genes in aortas between the two group. (H) Representative imaging results of contrast-enhanced computed tomography (CT) scanning of patients with aortic dissection, coronary angiography (CAG) of patients with acute myocardial infarction (AMI). The yellow and blue arrows indicate lesion sites in aortas and coronary arteries. (I) Clinical tests for serum biomarkers (Myoglobin, CK-MB, hsCRP and BNP) between AMI or AD patients. (Student’s t-test, \**P*≤ 0.05, \*\**P*≤ 0.01, \*\*\**P≤*0.001, ns, no significance.) (J) A corresponding ELISA kit developed by our team for detecting serum CTHRC1 levels. (K) CTHRC1 point-of-care testing (POCT) equipment, providing rapid and reliable results from a single drop of in less than 10 minutes. (L) The process of assessing chest pain during ambulance transport or in the ER.

## STAR★METHODS

### KEY RESOURCES TABLE

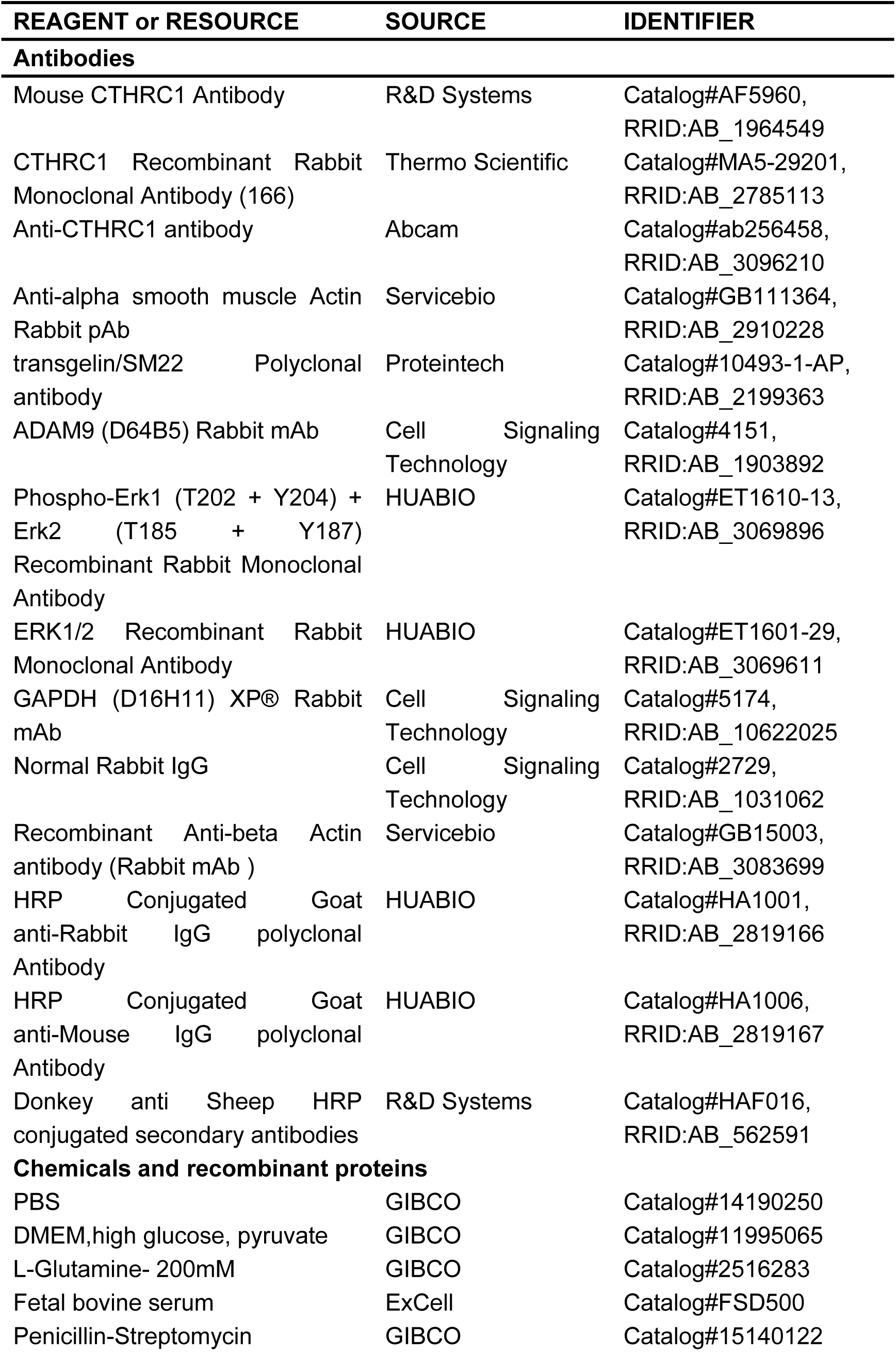
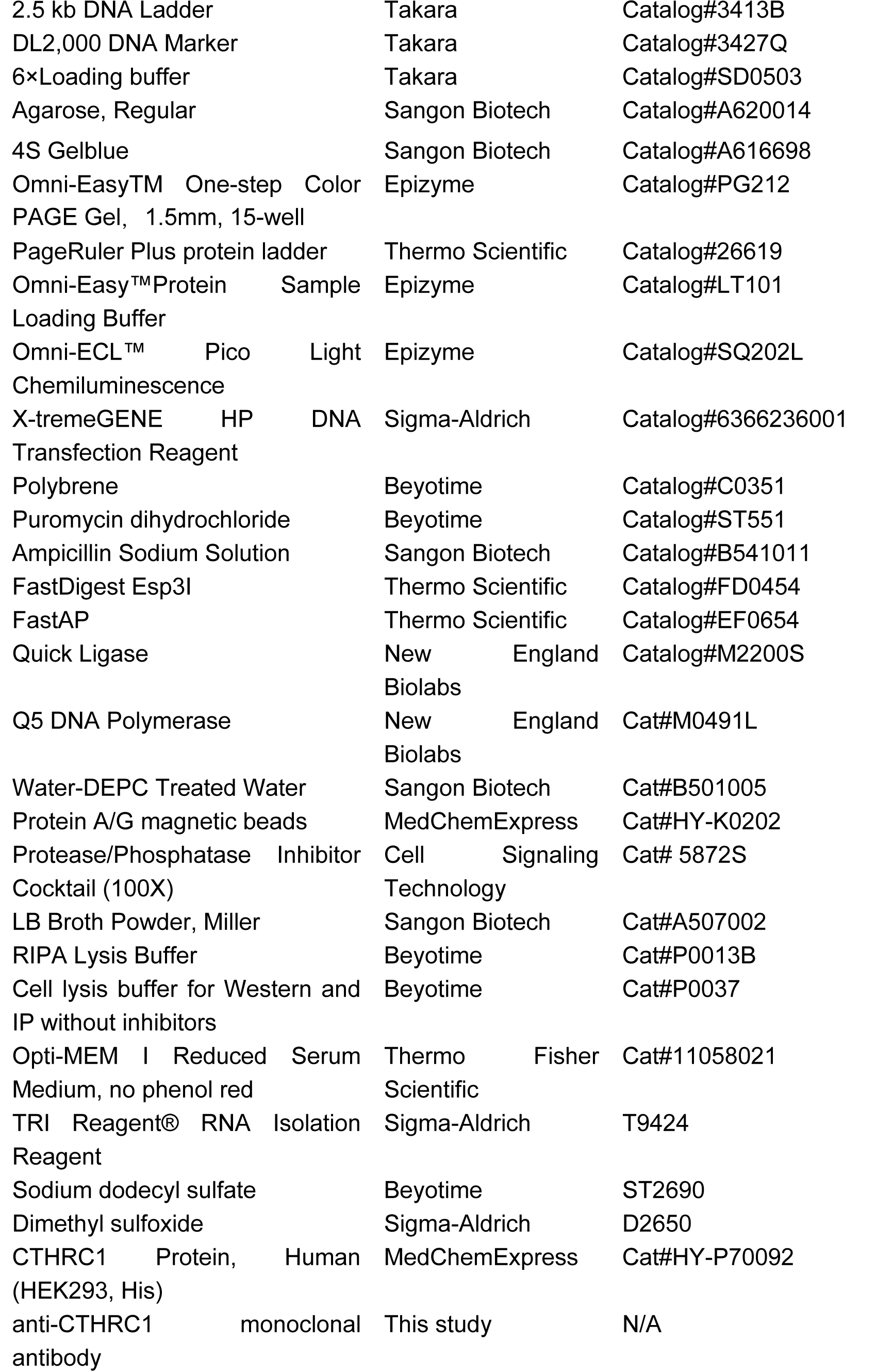
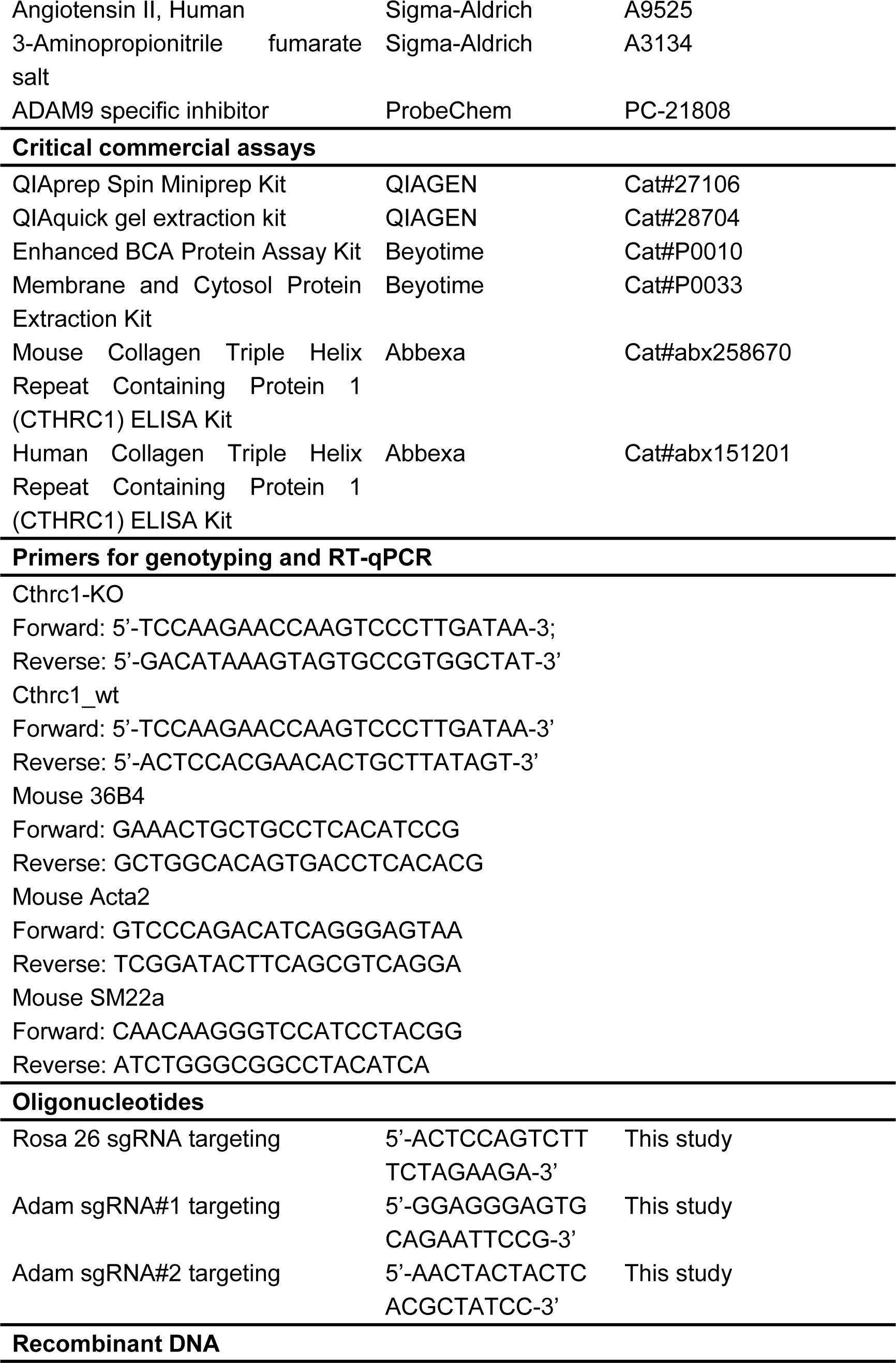
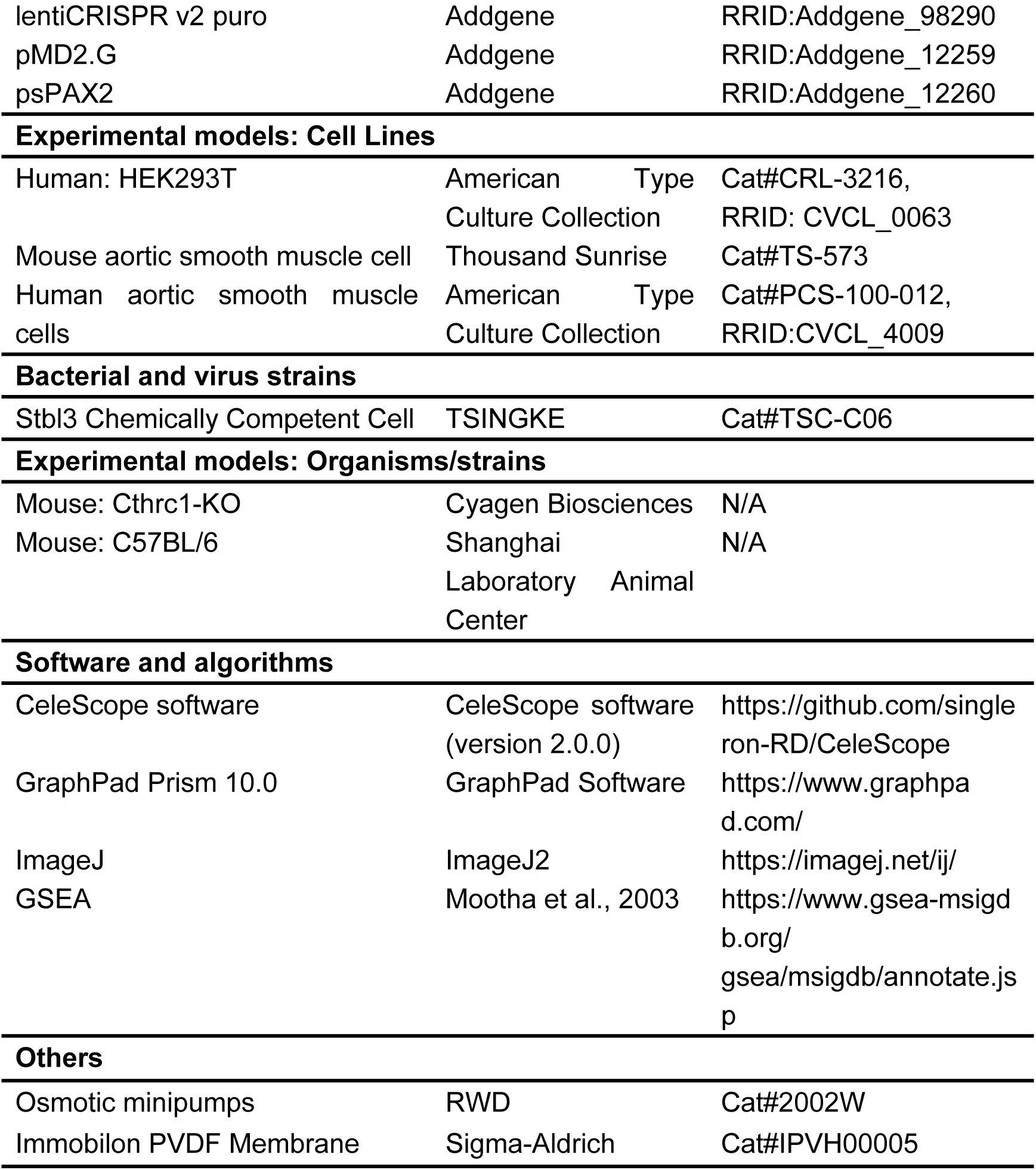

### RESOURCE AVAILABILITY

#### Lead contact

Further information and requests for resources (including code) should be directed to and will be fulfilled by the Lead Contact, Xiaoqing Wang.

#### Materials availability

Further information and requests for reagents may be directed to, and will be fulfilled by Prof. Xiaoqing Wang. A list of critical reagents (key resources) is included in the Key resources table. For additional materials, please email the lead contact for requests. Some material may require requests to collaborators and/or agreements with various entities. Material that can be shared will be released via a Material Transfer Agreement.

#### Data and code availability

The processed sequencing data in this paper have been deposited into the NCBI GEO database. Additional source data are available at Mendeley Data: https://data.mendeley.com/datasets/10.17632/g6zhdsc2tc.1.

### EXPERIMENTAL MODEL AND SUBJECT DETAILS

#### Animal Experiments

All animal protocols were approved by the National Institutes of Health Guide for the Care and Use of Laboratory Animals. All mice were fed normal laboratory diet, had free access to standard drinking water, and kept at approximately 21°C with 55% humidity and a 12-hour light/12-hour dark cycle. Six-to-eight week-old C57BL/6 male and female mice were purchased from Shanghai Laboratory Animal Center (SLAC). Mouse AD model was induced by simultaneous infusion of BAPN (150 mg/kg per day) and Ang-II (1000 ng/kg per min) using osmotic minipumps (RWD, 2002W) for 14 days. Cthrc1 knockout mice were constructed by Cyagen Biosciences. To monitor the structure and maximum diameter of blood vessel, the VEVO 3100 echography device (Fujifilm-Visual Sonics, Toronto, Canada) was used for obtaining the ultrasound images of the living mice. All mice were killed by pentobarbital overdose at the indicated time points to collect blood and tissue samples. Male and female mice were investigated similarly and independently.

#### Generation of *Cthrc1* knockout mice

*Cthrc1* KO mice were generated using CRISPR/Cas-mediated genome engineering. Two single-guide RNAs (sgRNAs) were designed to target the site of exon 2 and exon 3 in Cthrc1 genome, using the CRISPR design tool. The sgRNAs plasmids with Cas9 mRNA were co-injected into C57BL/6J mouse zygotes and surviving zygotes were transferred into pseudopregnants. The genotyping of Cthrc1 KO mice was confirmed by PCR amplification followed by DNA sequencing analysis. The tail tip of each mouse was obtained to extract genomic DNA. The targeting region was amplified by PCR genotyping primers. The sequences of primers were listed in supplementary tables.

#### Human subjects

Experimental procedures with human aortic tissues and blood samples were approved by the institutional review boards of the hospital. Informed consent was obtained for all recruited human participants. Ethical approval was consistent with the Declaration of Helsinki.

#### Cell culture

Mouse aortic smooth muscle cells (MOVAS, ATCC CRL-2797) and Human aortic smooth muscle cells (HASMC, ATCC PCS-100-012) were originally obtained from American Tissue Culture Collection Biobank (Manassas, VA, USA) and were maintained in DMEM (Gibco) with 10% FBS (FSD500, ExCell), penicillin/streptomycin 100 U/ml (15140122, Gibco) and 2 mM L-glutamine (25030081, Gibco), at 37°C with 5% CO2. And confluent MOVAS and HASMCs (80%–90%) were treated with recombinant human CTHRC1 Protein (1ug/mL) (HY-P70092, MedChemExpress) to investigate its effect on vascular remodeling in vitro. MOVAS and HASMCs within four-to-six passages was used in the following experiments.

### METHOD DETAILS

#### Single-cell capture, scRNA-seq library preparation and sequencing

Single-cell capture and library preparation were conducted using the Singleron Matrix® Single Cell Processing System. Fresh aortic tissues were preserved in the sCelLiveTM Tissue Preservation Solution (Singleron) on ice within half an hour. After being washed three times with Hanks Balanced Salt Solution (HBSS), the specimens were cut into small pieces and digested with 3 mL of sCelLiveTM Tissue Dissociation Solution (Singleron) using the Singleron PythoN™ Tissue Dissociation System at 37 °C for 15 minutes. The suspension of cells was collected and filtered with a 40-micron sterile strainer. Then, the GEXSCOPE® red blood cell lysis buffer (RCLB, Singleron) was added, and the mixture [Cell: RCLB=1:2 (volume ratio)] was incubated at room temperature for about 8 minutes. After centrifuging the mixture at 300 × g at 4℃ for 5 minutes to eliminate the supernatant, it was softly resuspended in PBS (HyClone). Single-cell suspensions (2×10^5^ cells/mL) were subsequently loaded onto a microwell chip using the Singleron Matrix® Single Cell Processing System. After collecting the Barcoding Beads from the above microwell chip, the captured mRNA undergoes reverse transcription to produce cDNA, which is then amplified using PCR. Following amplification, the cDNA is broken into fragments and linked with sequencing adapters. The scRNA-seq libraries were developed according to the GEXSCOPE® Single Cell RNA Library Kits (Singleron) guidelines. The libraries were then diluted to a concentration of 4 nM, combined, and sequenced using the Illumina NovaSeq 6000 platform with 150 bp paired-end reads.

#### Bulk RNA-seq

High-throughput mRNA sequencing was performed by the LC-Bio Technology co.ltd., (HangZhou, China). The quality of total RNAs was assessed by BioAnalyzer 2100 (Agilent Technologies). RNA libraries were constructed with TruSeq Stranded Total RNA Library Prep kit (Illumina). Libraries were quality-controlled for size distribution and quantified with Qubit Fluorometer (Thermo Fisher Scientific). The libraries were sequenced on an Illumina HiSeq sequencer for 150-bp paired-end reads according to the manufacturer’s instructions.

#### Histological staining

After intraperitoneally injected at overdose pentobarbital, mice were completely infused by PBS from the heart. Then aortic tissues were isolated and fixed in 4% paraformaldehyde (PFA) overnight followed by embedded in paraffin. Aorta sections (4 μm thick) were stained with hematoxylin and eosin (H&E), Verhoeff elastic–van Gieson (EVG) and Alcian blue for histopathological analyses. H&E, EVG and Alcian blue staining were performed according to the manufacturer’s protocol. Images were acquired using the Pannoramic 250 FLASH scanner (3DHISTECH, Budapest) and processed by CaseViewer2.4 software (3DHISTECH, Budapest). For good presentation, the pictures were arranged with Photoshop and Illustrator (Adobe, 2024) according to the guidelines of this journal.

#### Immunohistochemical staining and Immunofluorescence staining

For immunohistochemical staining, the aorta sections (4 μm thick) in paraffin-embedded were dewaxing and antigen retrieval with a Tris-EDTA (pH 9.0), followed by treatment with 3% hydrogen peroxide. Then tissue sections were permeabilized with 0.3% triton and blocked with 5% normal donkey serum for 30 min at 37℃. Slides were incubated with Mouse CTHRC1 primary antibody (AF5960, R&D System) at 4°C overnight, then HRP-conjugated anti-Sheep IgG secondary antibody (HAF016, R&D System) were added the next day at room temperature for 1 hour. Lastly, tissues were stained with a DAB kit.

Paraffin-embedded aortic tissues were deparaffinized and rehydrated. Secondly, the aorta sections were repaired with EDTA antigen repair solution (pH 8.0) and blocked with 3% BSA and incubated overnight at 4°C with the α-SMA, SM22α primary antibodies (GB111364-100, Servicebio; 10493-1-AP, Proteintech). After several washes with PBS, the aorta sections were incubated with fluorescent secondary antibody Goat Anti-Rabbit IgG (CY3) (GB21303, Servicebio) for 50 minutes at room temperature. Lastly, DAPI solution (G1012, Servicebio) was dripped into the circle and incubated at room temperature for 10 min in the dark. Immunofluorescence was visualized using a confocal microscope (Nikon Eclipse C1, Japan) and scanned by Pannoramic MIDI (3DHISTECH, Budapest). For each image, the median fluorescence intensity was quantified by Image J software.

#### Quantification of plasma level of CTHRC1

Mouse plasma was collected by centrifugation of blood from eyeballs (3000×rpm, 20 min). Then the secreted CTHRC1 was measured using Mouse Collagen Triple Helix Repeat Containing Protein 1 (CTHRC1) ELISA Kit (abx258670, Abbexa) according to the manufacturer’s instructions. Similarly, serum CTHRC1 levels were quantified in patients and controls by collecting 5 ml venous blood into plain tubes and subjected to extensive centrifugation to isolate the serum. Additionally, the subsequent measurement of serum CTHRC1 levels was conducted using Human Collagen Triple Helix Repeat Containing Protein 1 (CTHRC1) ELISA Kit (abx151201, Abbexa), following the same protocol as previously described.

#### CTHRC1 monoclonal antibody production

His-tag fusion proteins of CTHRC1 (encoding 31-243 amino acids of human CTHRC1) was constructed into plasmid, expressed and purified according to manufacturer’s protocols. The purified human recombinant CTHRC1 protein was then used as the antigen to immunize mice for generation of anti-CTHRC1 monoclonal antibodies. Immunized mice meeting the serum titer criteria undergone the subsequent fusion with myeloma cells for ELISA assay to screen monoclonal antibodies (Huabio Biotechnology Co., Ltd). An monoclonal antibody was produced and labeled as 1A5-10, the immunoglobulin subtype of which was IgG2b. The purity of this monoclonal antibody was 90%, and the concentration was 1mg/ml.

#### Generation and transfection of lentivirus plasmids

To knockout ADAM9 expression in cells, two independent guide sequences were designed targeting Adam9 gene. The sequence of the two Adam9 sgRNA are: 5’-GGAGGGAGTGCAGAATTCCG-3’, 5’ -AACTACTACTCACGCTATCC-3’. Each pair of oligonucleotides were annealed and ligated into the lentiCRISPR v2 vector (#52961; Addgene). Lentiviruses were produced in HEK293T cells with 90% confluency. The following day cells were co-transfected using 50 μl X-TremeGene Transfection Reagent (Roche) diluted in 0.5 ml of Opti-MEM (ThermoFisher Scientific) that was combined with 2 μg of the plasmid pool DNA and a mixture of 1.5 μg psPAX2 and 0.6 μg pMD2.G. After 12 h, the culture medium was refreshed. At 72 h post-transfection, supernatant-containing virus was collected and passed through a 0.45-μm filter. Virus was harvested, aliquoted, and used to transfect mouse aortic smooth muscle cells in a 6-well-plate.

#### In vitro assays incubated with recombinant CTHRC1 (rhCTHRC1)

To validate the effect of CTHRC1 on contractile-to-synthetic phenotype switching in vitro, we treated aortic smooth muscle cells with 1 μg rhCTHRC1 for 0, 1, 3, 6, 9, 12, 24h. Then the cells were washed with ice-cold PBS and subjected to the following experiments of bulk RNA-seq, immunoblotting and RT-qPCR analysis.

#### Protein extraction and western blotting experiments

Total protein and membrane fractions were extracted by using RIPA lysis Buffer (P0013B, Beyotime) and Membrane and Cytosol Protein Extraction Kit (P0033, Beyotime) respectively. For western blotting experiment, equivalent extracted protein samples were resolved by 7.5% or 10% sodium dodecyl sulfate polyacrylamide gel electrophoresis and transferred onto polyvinylidene fluoride membranes (PVDF). Then membranes were blocked (5% skim milk in TBST buffer) at room temperature for 1h. Subsquently, the membranes were incubated with primary antibody overnight at 4℃. After washed with TBST buffer for 3 times, membranes were incubated with secondary antibody at room temperature for 1h and visualized by an electrochemiluminescence (ECL) system. The primary antibody used were as follows: anti-CTHRC1 at 1:1000 (ab256458, Abcam); anti-α-SMA at 1:1000 (GB111364, Servicebio); anti-SM22α at 1:1000 (10493-1-AP, Proteintech); anti-ADAM9 at 1:1000 (#4151, CST); anti-p-ERK1/2 at 1:1000 (ET1610-13, HUABIO); anti-ERK1/2 at 1:1000 (ET1601-29, HUABIO); anti-GAPDH at 1:5000 (#5174, CST); anti-β-actin at 1:1000 (GB15003, Servicebio); anti-Na/K-ATPase at 1:5000 (T55159, Abmart). Evidently, protein extraction from tissues was similar as above.

#### Co-Immunoprecipitation and mass spectrometry

The protein of membrane fractions from vascular smooth muscle cells was extracted using Membrane and Cytosol Protein Extraction Kit (P0033, Beyotime). A BCA assay (Beyotime, P0010) was utilized to measure protein membrane fraction concentration. Then the total amount of 800 μg protein was dissolved in lysis buffer and divided into IP : CTHRC1 (ab256458, Abcam) and normal rabbit IgG groups (#2729, CST) followed by shaking overnight at 4°C. The lysate were incubated another 2 hours on a rotating mixer with protein A/G magnetic beads (HY-K0202, MedChemExpress) the next day and then washed with PBST buffer for 4 times. Lastly, the mixture was eluted in lysis buffer for further western blotting or mass spectrometry. After excising the SDS-PAGE gels, enzyme digestion was performed, and the peptide fragments were desalted through chromatography. The desalted peptides were dried for further processing. Subsequently, they were processed through an ESI source and tandem mass spectrometry (MS/MS) using an Orbitrap Fusion Lumos Tribrid (Thermo, USA) that was online with the HPLC. Protein identification was carried out using Proteome Discoverer 2.1 software by searching the UniProt Consortium.

#### Quantitative real-time polymerase chain reaction

According to standard guidelines, total RNAs from aortic tissues or cultured vascular smooth muscle cells were extracted with TRIzol reagent (T9424, Sigma-Aldrich). The RNA concentrations and purity were determined using a UV/VIS Nano Spectrophotometer (Cobra, Oxford). Subsequently, a amount of 1 μg mRNA was transformed to cDNA utilizing the HiScript II Reverse Transcriptase Reagent kit with gDNA eraser (Vazyme, China). Quantitative RT-PCR reactions were conducted using the AceQ Universal SYBR qPCR Master Mix (Vazyme, China) on a LightCycler® 96 Instrument (Roche, USA). The primers are displayed in supplementary tables.

### QUANTIFICATION AND STATISTICAL ANALYSIS

#### Analysis of scRNA-seq data

Gene expression matrices were generated using the CeleScope software (version 2.0.0) (https://github.com/singleron-RD/CeleScope), which is a collection of bioinformatics analysis pipelines developed by Singleron for processing single cell sequencing data. Then the expression matrix files were imported into Python (version 3.9.18), and the Scanpy package (version 1.9.6)^45^ was used for downstream analysis. We performed the following quality control steps: (i) genes expressed in less than 10 cells were removed; (ii) cells that expressed fewer than 500 genes were excluded; (iii) cells with over 10% of unique molecular identifiers (UMIs) derived from mitochondria were not considered. After removing low-quality cells, the count data were normalized using the pp.normalize_total and pp.log1p function as implemented in the Scanpy package. The scanpy.pp.highly_variable_genes function with batch_key parameter was then used to identify the highly variable genes (HVGs). Subsequently, principal component analysis (PCA) was performed using the scanpy.tl.pca function to map these HVGs to the low-dimensional subspace, and the Harmony algorithm^96^ was utilized to remove batch effects with the harmonypy.run_harmony function. After using the scanpy.pp.neighbors function to compute the neighborhood graph of cells, the scanpy.tl.leiden function with resolution of 0.5 was utilized to identify cell clusters. To visualize the clustered cells, we first embedded the neighborhood graph into two dimensions using the scanpy.tl.umap function, and then used the scanpy.pl.umap function to generate the scatter plot in uniform manifold approximation and projection (UMAP) basis. Each cell cluster was annotated based on the known marker genes. Heatmaps showing expression level of selected genes in the different cell types were generated using scanpy.pl.matrixplot function. To identify differentially expressed genes (DEGs) for the certain cell type between different conditions, we used scanpy.tl.rank_genes_groups function with default parameters. Genes with the threshold of fold change >2 and false discovery rate (FDR) <0.05 were considered as DEGs. Gene Ontology (GO) analysis was performed on DEGs using the R package clusterProfiler (version 4.6.2).^97^ To evaluate and compare cell-cell communications between different cell types or conditions, we used the R package CellChat (version 2.1.2)^59^ that integrates gene expression with the known interactions between signaling ligands, receptors and cofactors.

#### Data analysis of RNA-seq

Pair-end FASTQ files were mapped to the Mus_musculus.GRCm38.101 and Homo_sapiens.GRCh38.112 reference genomes respectively, via using the HISAT2 package. Then, the FPKM values were calculated with StringTie and Ballgown (http://www.bioconductor.org/packages/release/bioc/html/ballgown.html). Differential gene expression was analyzed using DESeq2, with genes having a *P* value ≤ 0.05 and absolute fold change ≥ 2 recognized as differentially expressed genes (DEGs). Heatmaps were generated, and Gene Ontology (GO) and KEGG pathway analyses were performed on DEGs using the OmicStudio tools at https://www.omicstudio.cn/tool. Gene set enrichment analysis (GSEA) was conducted using GSEA (v4.1.0) and MSigDB to identify gene sets involved in specific pathways.

#### Molecular modeling analysis of CTHRC1-anti-Cthrc1 Ab and CTHRC1-ADAM9 interactions

The highly integrated HDOCK server was used for protein–protein docking (http://hdock.phys.hust.edu.cn/) based on a hybrid strategy. Briefly, the protein structures were created for CTHRC1, anti-Cthrc1 Ab and ADAM9 by AlphaFold3 server (https://alphafold3.org/). MM/GBSA algorithm of HawkDOCK SERVER (http://cadd.zju.edu.cn/hawkdock/) was employed to calculate the atomic affinity grid under the condition of docking simulation. CTHRC1-anti-Cthrc1 Ab and CTHRC1-ADAM9 interactions were selected based on criteria of interacting energy and geometrical matching quality by calculating Docking Score. Finally, we used Pymol 2.4 software to visualize these hydrophobic interaction sites of CTHRC1-anti-Cthrc1 Ab and CTHRC1-ADAM9.

#### Statistical analysis

Statistical analysis was performed in Prism 10 (GraphPad Software). Kaplan-Meier survival curves were adopted to assess survival rates of mouse, and the differences between groups were analyzed by using the Log-rank (Mantel-Cox) test. The AD or NA incidence was analyzed using the Chi-square test. All continuous data are presented as mean ± SEM. For comparison between two groups, an unpaired two-tailed Student’s t-test was used assuming that the data have the equal variance. Otherwise, t test with Welch’s correction was performed. Similarly, when data were compared among more than 3 groups, a ordinary one-way ANOVA was used following by Bonferroni’s multiple comparisons test based on the equal variance. Providing that there were significantly equal variance among more than 3 groups, Brown-Forsythe and a Welch ANOVA test was performed, using the Dunnett’s T3 method. The threshold of statistical significance was *P* ≤ 0.05.

