## Supplementary figures and images for "CTHRC1 is a new therapeutic target and serum diagnostic biomarker for aortic dissection"

### Supplemental_figure_1

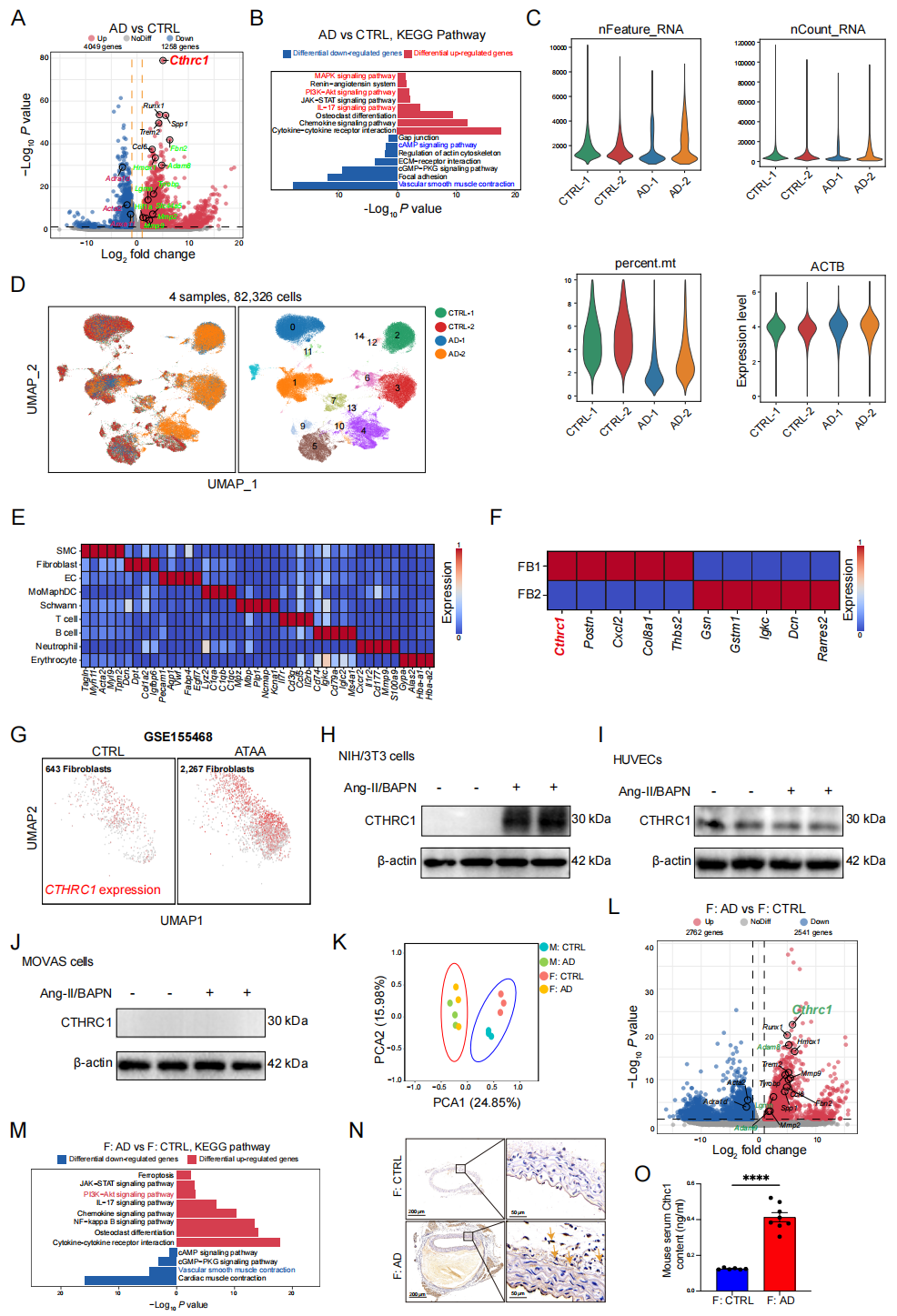

### Supplemental_figure_2

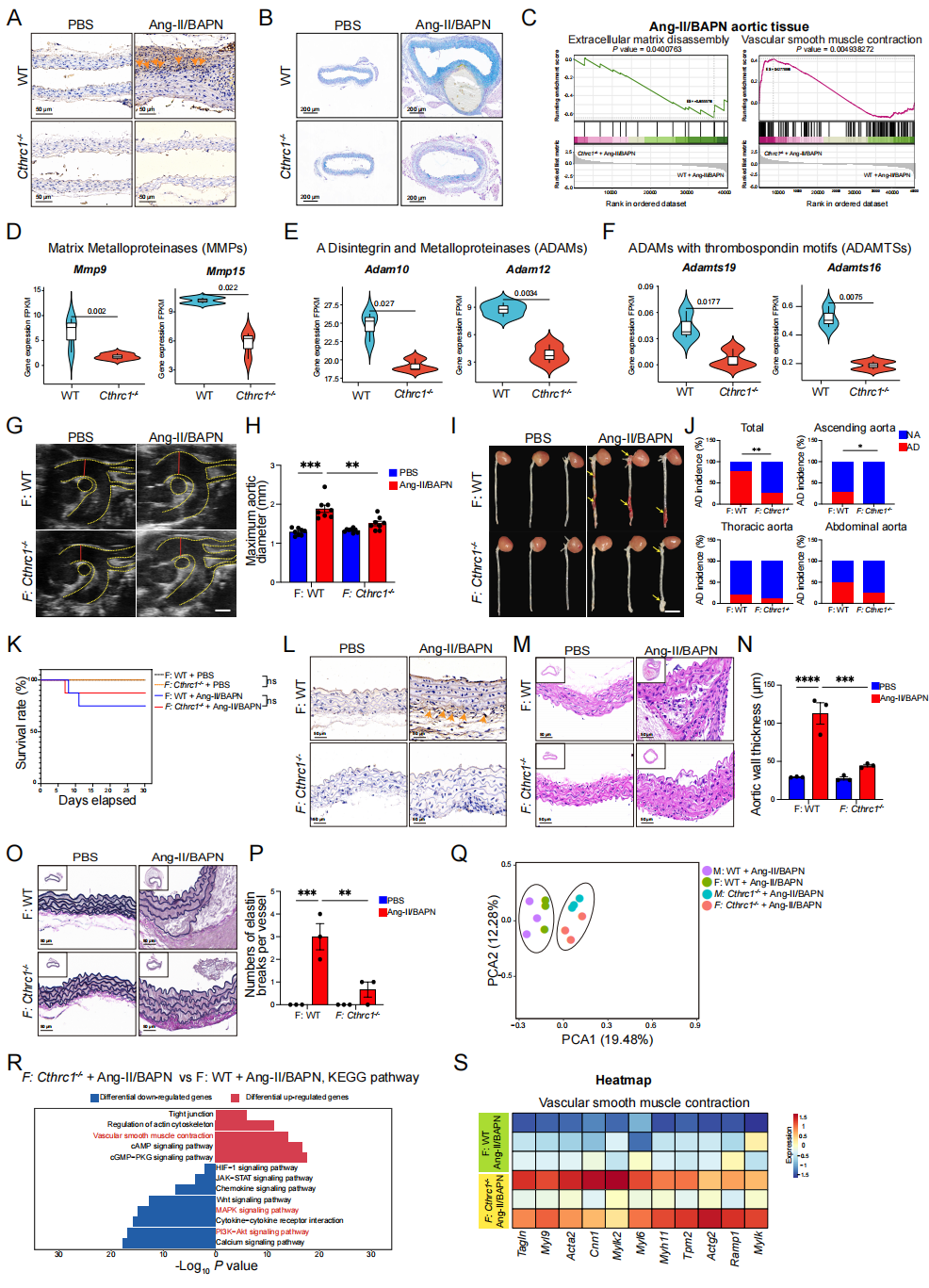

### Supplemental_figure_3

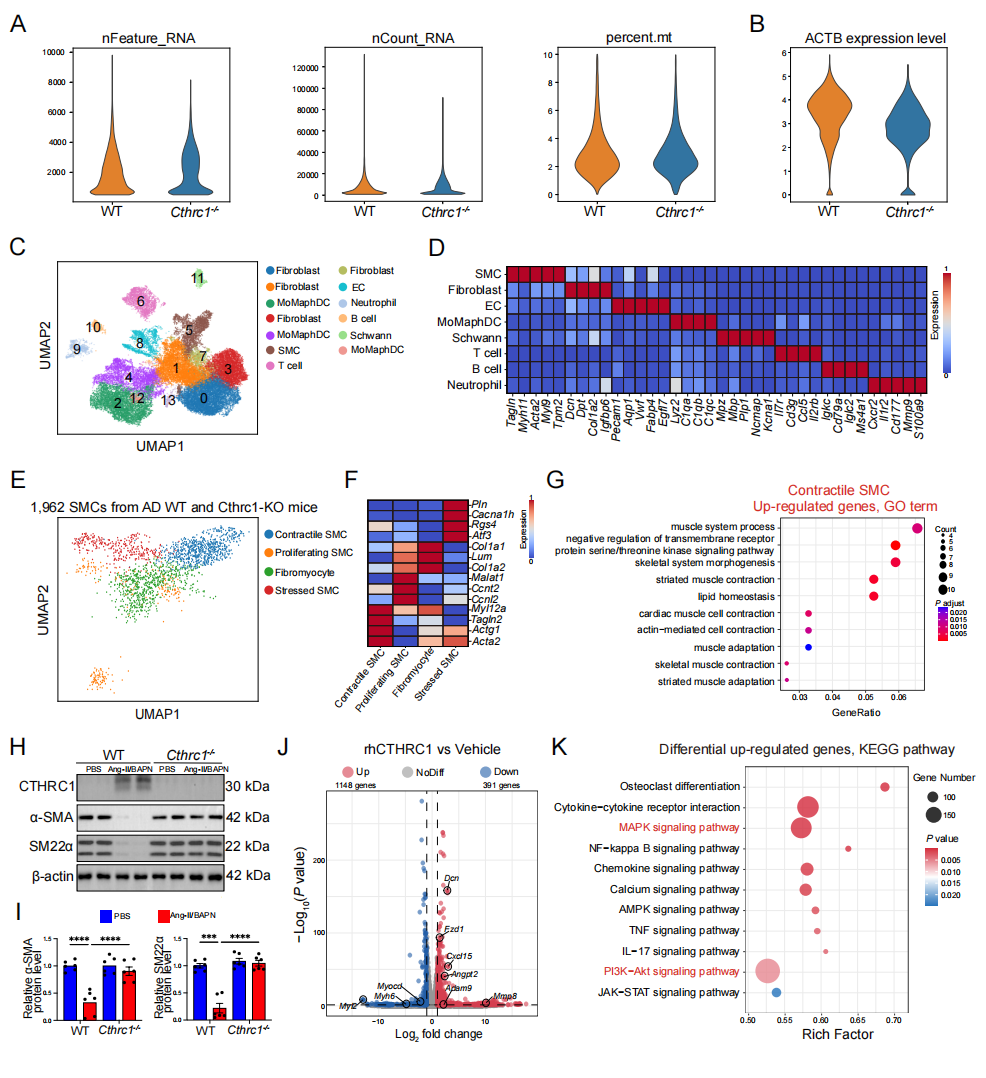

### Supplemental_figure_4

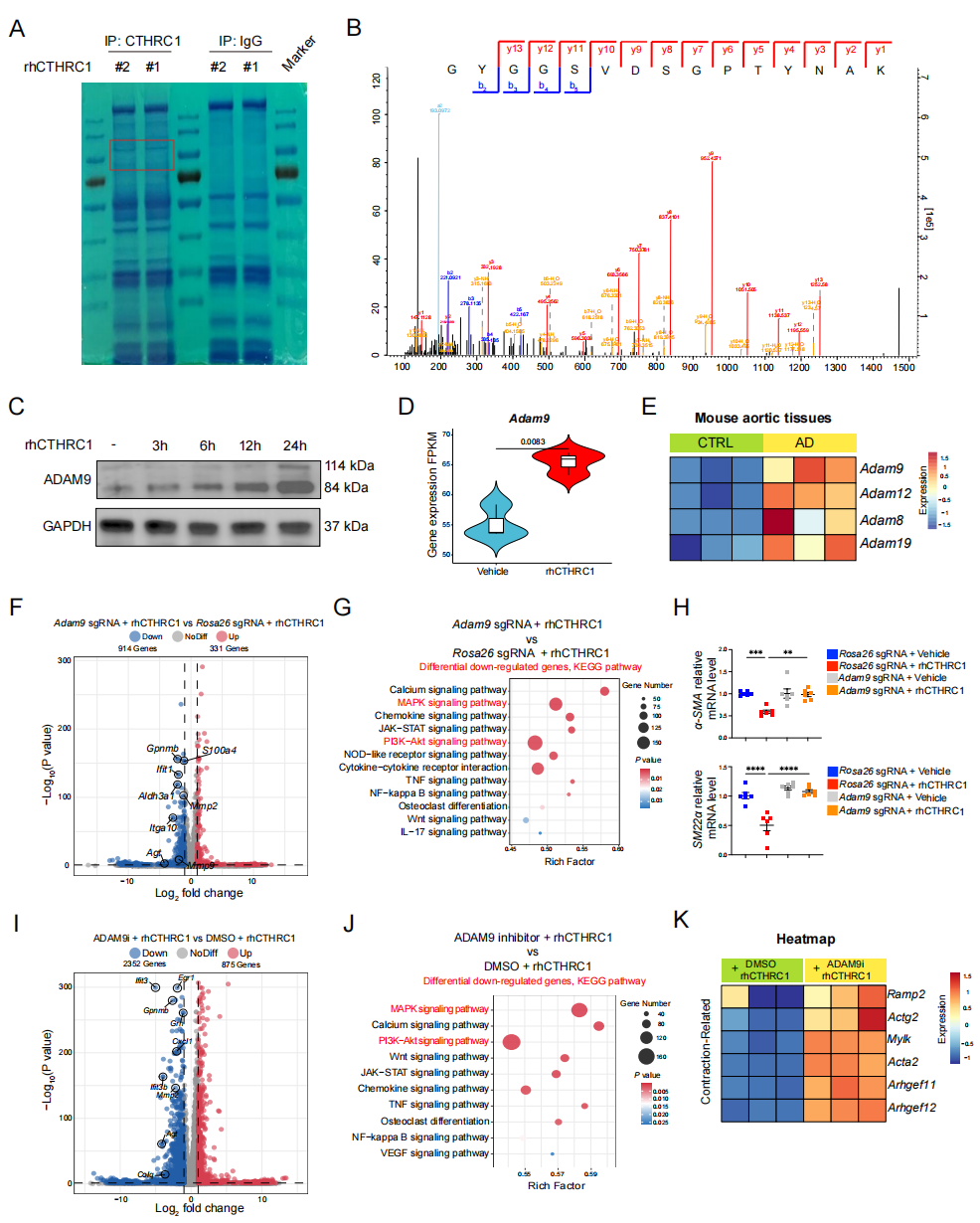

### Supplemental_figure_5

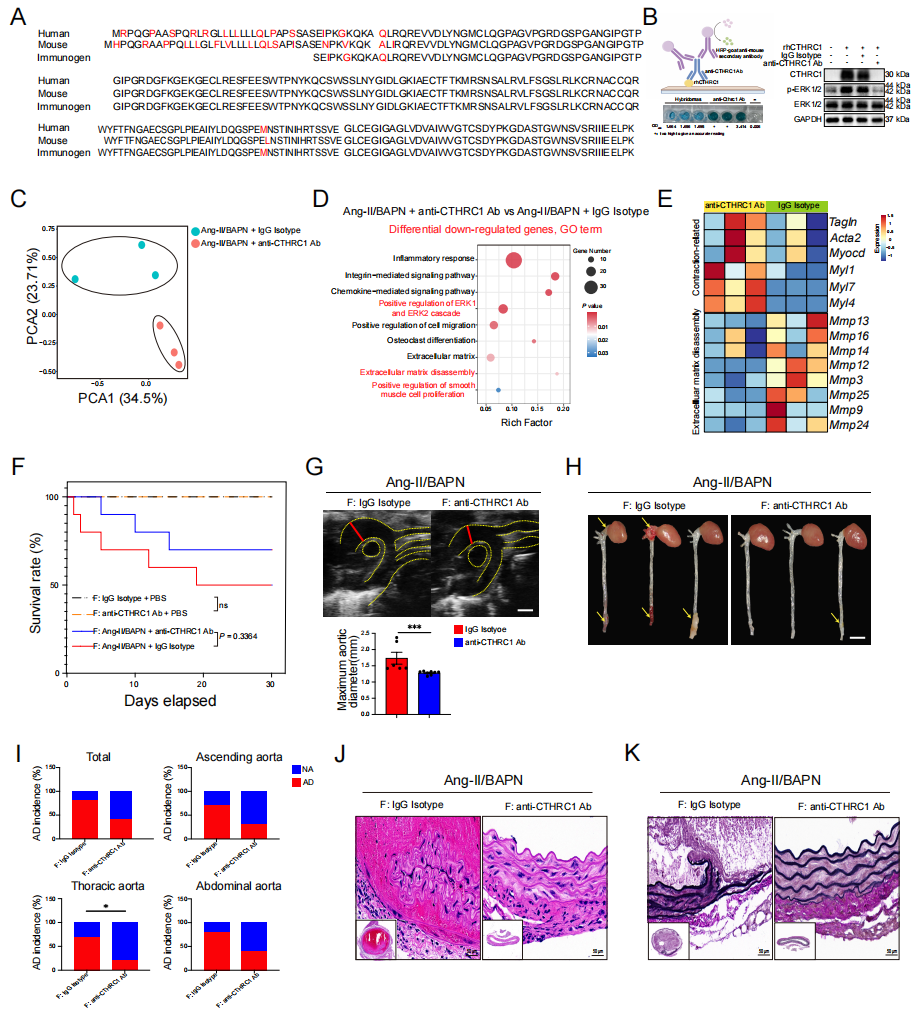

### Supplemental_figure_6

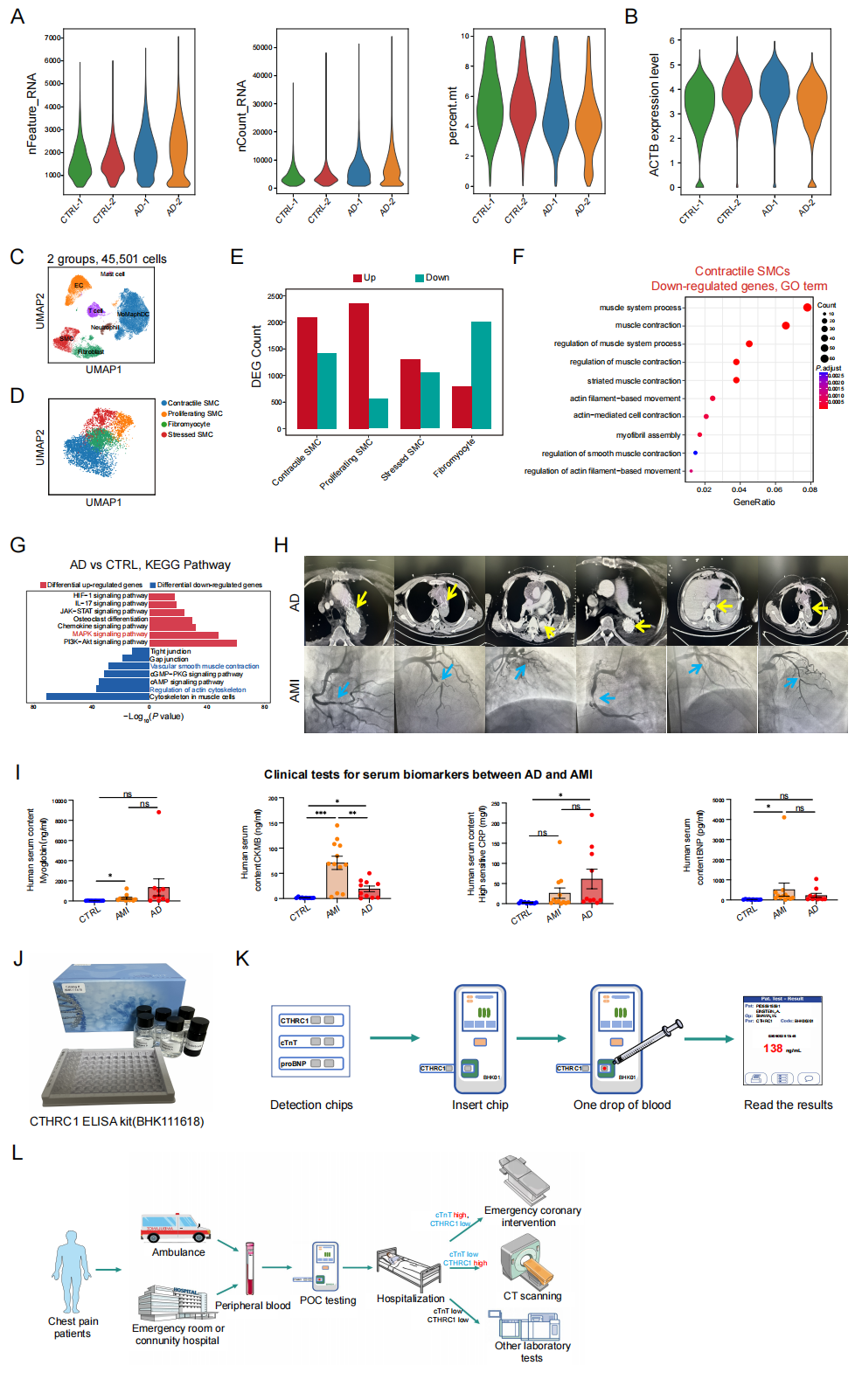
